# Moirai: single-cell trajectory inference grounded in gene-level expression dynamics

**DOI:** 10.64898/2026.08.05.742709

**Authors:** Amber H.B. Fijn, Gustavo S. Jeuken

## Abstract

Underlying the development of multicellular organisms is the process of cell differentiation, which is governed by the concerted and sequential change in gene expression. Various methods have been developed that employ scRNA-seq data to infer the position of a cell along a pseudo-temporal axis and identify relevant genes involved in the process. These trajectory inference methods typically rely on global transcriptomic changes and mathematical methods. However, overemphasis on large-scale transcriptomic changes may impair sensitivity to identify branching points and convergent trajectories, which are rather governed by small-scale transcriptional events. Motivated by this, we developed *Moirai*, a graph-based trajectory inference method that identifies gene expression patterns that change dynamically over a developmental continuum and leverages these to define a common pseudotime axis between all cells. In doing so, *Moirai* shifts the focus to individual gene dynamics, which enhances its ability to detect putative branching points that are masked by global transcriptomic similarities. We apply *Moirai* to four developmental datasets, where we demonstrate its ability to recover gene expression patterns of genes with a known involvement in the respective developmental process, motivating their use for defining a cell’s pseudotime. We furthermore show that *Moirai* can robustly infer gene expression patterns across different embedding approaches, highlighting the value of moving the focus of the inference process to the small-scale transcriptional dynamics.

## Introduction

Cell differentiation is a fundamental process underlying the development of multicellular organisms. It continues throughout an organism’s life and is vital for the maintenance and repair of various adult tissues. Driving the process of differentiation is the sequential and concerted change in gene expression, organized by the gene regulatory programs acting in response to intra- and extracellular stimuli. Together, these programs effectively drive a pluripotent, undifferentiated cell towards a more specialized cell state. Development towards this specialized cell state was long thought to consist of a sequence of well-defined intermediate states. However, this idea has changed, shifting due to a recognition of the continuous nature of cell differentiation, where the paths towards a specialized cell state appear less well defined than long believed [1, 2]. Illustrating this point, various studies have already revealed the occurrence of terminal state convergence, where cells with distinct molecular origins converge on similar end states, making the developmental history of a terminally differentiated cell ambiguous [3, 4, 5, 6].

To study the cellular events that characterize the developmental path of a cell requires data reflecting a cell population that spans this continuum. Single-cell RNA sequencing (scRNA-seq) data capture the full transcriptomic state of a population of cells and thus grant the opportunity to infer their developmental stage. This is particularly true for samples taken from homeostatic adult tissues, where the continuous, asynchronous turnover translates into a cell population covering many (if not all) stages of differentiation [7]. Since these data represent a snapshot in time, the challenge arises to infer the shape of the developmental topology and the position of cells along it using only their transcriptional profile as guides. In doing so, transcriptional dynamics may be studied over pseudo-time, which can provide insights into differentiation and the role of various molecular agents. Complicating this feat is the inherent noisiness of scRNA-seq data, arising from both technical and biological sources [8, 9]. This may conceal the developmental signal, which should be identified for meaningful trajectory inference. These challenges underscore that computational trajectory inference requires careful design.

Several trajectory inference (TI) methods have been developed that exploit cell-to-cell similarity to infer a relative ordering of cells along a pseudo-temporal axis. These methods assume that scRNA-seq data reflect the full developmental continuum and that this continuum can be summarized into a two-dimensional trajectory. To capture this trajectory, TI methods generally make use of either minimal-spanning tree (MST) or nonlinear embedding approaches [10]. MSTs are acyclic graphs in which all nodes are connected so that the total edge weight is minimized [11]. This effectively connects all nodes in such a way that only a single path exists between them. In TI applications, the nodes of the graph represent (clusters) of cells, and the edge weights reflect some similarity measure (e.g., covariance) between connected clusters [12]. Lineages are then defined as the ordered set of clusters visited along paths over the MST. MSTs successfully derive global structures from large datasets [13], but have shown to be unstable [10]. Non-linear embedding methods represent the developmental data in lower dimensional spaces, from which the trajectory is inferred using various approaches. For instance, Wishbone uses PCA and diffusion map to obtain an embedding of the data from which a *k*-nearest neighbor graph is constructed [14]. By repeatedly sub-sampling the edges and computing the shortest-path distance between cells, Wishbone is able to identify bifurcations and map cells to a trajectory.

Most TI methods depend on global transcriptomic changes to derive pseudotemporal ordering of the cells. Yet, increasing evidence suggests that just a subset of genes are sufficient to mark a cell’s developmental fate [15, 16, 17]. These master regulators are highly connected genes that can switch between on/off states through positive feedback loops, thus robustly and decisively controlling the developmental path of a cell [16]. To illustrate, during hematopoiesis cross-antagonism between GATA1 and PU.1 effectively differentiates between the erythroid and myeloid lineage [17]. Although the changes induced by such regulators manifest on a transcriptome-wide scale, this occurs with a temporal delay. Overemphasis on global transcriptomic changes may then impact the ability of the model to capture developmental branching events which may limit interpretation gained from these models.

Recognizing the value of shifting the focus to the small-scale transcriptional changes, we developed *Moirai*, a graph-based trajectory inference method that unlike other methods leverages the individual expression patterns sampled across the developmental continuum to order cells temporally. Instead of relying only on global cell-to-cell similarity, *Moirai* identifies the genes that change dynamically across the graph and uses these dynamics to define a temporal axis common to all cells. In doing this, *Moirai* robustly estimates a cell’s pseudo-time and simultaneously captures the expression patterns that govern cell fate specification. Applying *Moirai* on four developmental datasets, we demonstrate *Moirai*’s ability to effectively identify developmentally relevant gene expression patterns and infer pseudotime using these. Spatial bias of the gene dynamics highlighted potential branching points between lineages, which were not reflected in the topology of the embedding. Moreover, with *Moirai* we identified a collection of genes across the four datasets without a known involvement in the respective developmental processes, further demonstrating its use for biological discovery.

## Methods

Motivated by the small-scale transcriptional changes that underlie key developmental events we set out to develop *Moirai*, a graph-based method that utilizes gene expression patterns sampled across the developmental continuum to predict pseudotime. Underlying *Moirai* are two key assumptions, which are (1) the expression of (a subset of) genes involved in cell fate specification changes gradually across pseudotime, and (2) these expression patterns can be captured across a population of cells. To capture these expression patterns, we embed single cells in a lower dimensional space which is used to construct a graph that captures the developmental topology. Here, we perform random walks from among which we identify expression patterns that change dynamically across the graph. We assume that these expression patterns belong to genes involved in cell fate specification. By defining a common pseudotime axis based on the expression patterns sampled across the graph, we finally infer pseudotime for all cells represented in the data.

The general workflow of *Moirai* comprises seven main steps (Figure 1):

- Step 1 – Build a k-NN graph.
- Step 2 – Construct a transition matrix.
- Step 3 – Perform random walks on the k-NN graph.
- Step 4 – Score gene dynamics sampled along the random walks.
- Step 5 – Select smooth gene dynamics and align random walks to identify common expression patterns.
- Step 6 – Update cell-cell transitions to bias walks towards smooth gene dynamics and repeat Step 3 - Step 6 for *i* iterations.
- Step 7 – Predict pseudotime from the aligned random walks.

**Figure 1:**
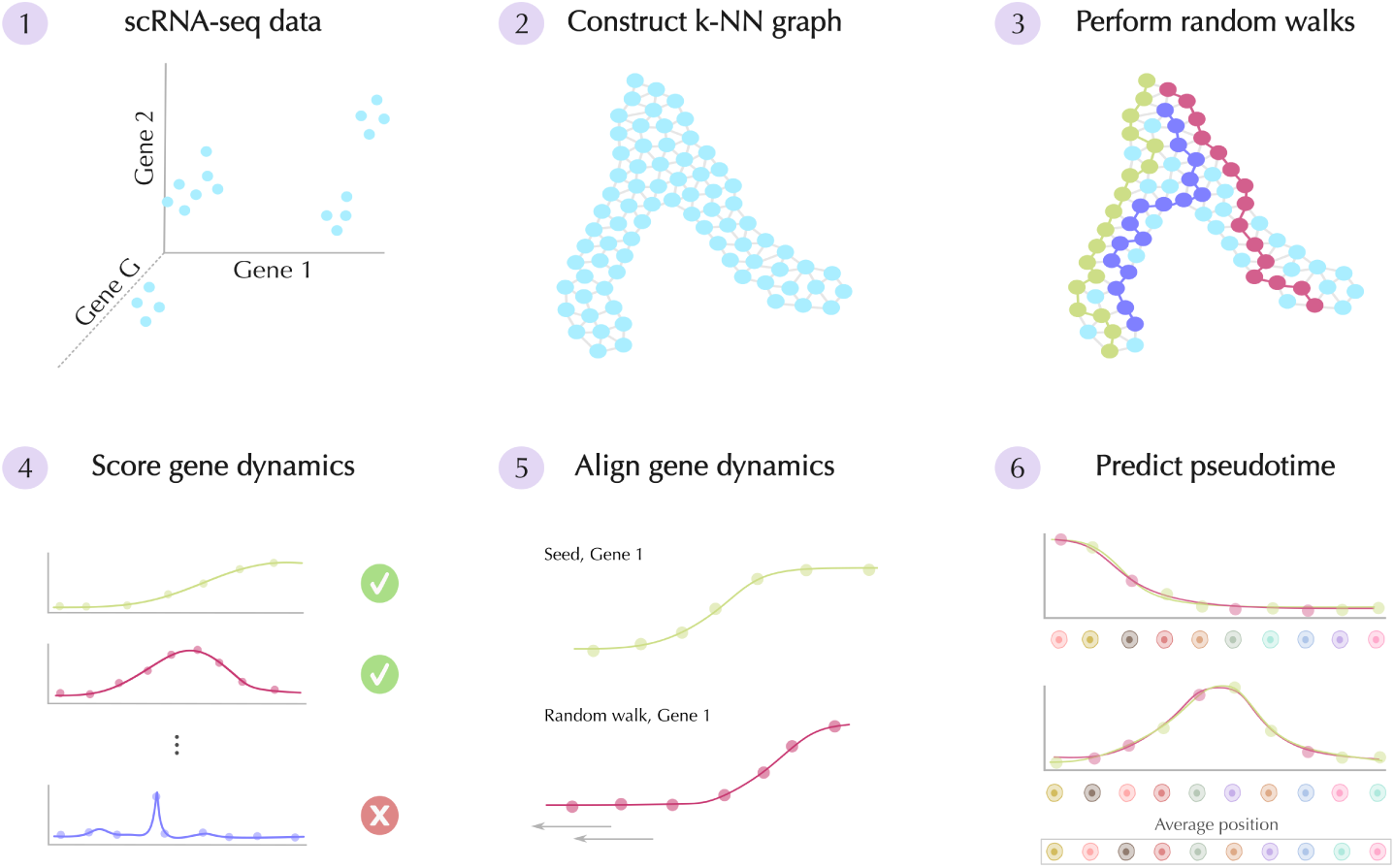
Overview of *Moirai*. *Moirai* predicts pseudotime by identifying those gene expression patterns that change dynamically over the developmental continuum, and leverages these to infer a cell’s position along this continuum. To do this, *Moirai* proceeds through six global steps. Single-cell RNA seq (scRNA-seq) data are provided to *Moirai* (1) and are embedded in a lower dimensional space on which a *k*-nearest neighbors (k-NN) graph is constructed (2). From the k-NN graph we obtain a transition matrix, which is used to perform random walks across the graph (3). These random walks are then scored based on the smoothness of the gene expression patterns they captured over the cells visited in the respective walk (4). We then compare those smooth expression patterns to see if similar dynamics were captured across different random walks (5). This is done by defining a representative random walk, a seed, to which all other walks are compared that captured a smooth expression pattern for the respective gene. If the two expression patterns can be aligned, the relative positioning is stored; effectively creating a pseudo-ordering for the cell populations captured in the respective random walks. Aggregating all alignments from Step 5, an average position can be predicted for a cell, which will then be scaled to obtain its final pseudotime (6).

### Step 1. Build a k-NN graph

To perform random walks across the developmental continuum, the data must be summarized in graph structure. Raw scRNA-seq data are log-normalized and highly variable genes are selected to reduce computational costs and limit the search space to those genes that vary in expression magnitude. We assume that among these genes are those involved in cell differentiation, which thus should impact the embedding most strongly. Taking the pre-processed data, we embed the cells in a lower dimensional space that is assumed to represent the global developmental topology which may be obscured by biological and technical noise in the high-dimensional representation. Dimensionality reduction is furthermore necessary to increase efficiency and accuracy of downstream graph construction. Various dimensionality reduction methods can be used and are supported by *Moirai*. By default, we compute a UMAP as it is suitable for studying developmental trends due to its ability to capture global relations between cells [18, 19]. The UMAP is constructed by first computing the PCA, from which we keep the first *z* components that are then used to compute the UMAP embedding.

Taking the cell’s coordinates in the embedded space, we build the *k*-nearest neighbor (k-NN) graph. Here, the nodes (*v*) represent the cells, and the edges (*e*) reflect the Euclidean distance between cells in the embedded space. The value of *k* depends on the sampling density of the cell types represented in the data. If relatively few cells of a particular cell type are captured in the data, their transcriptomic profiles may be expected to differ more strongly. To still capture smooth expression patterns, it becomes more important that cells connections are only drawn between the most similar pairs, which requires a lower *k*. At higher sampling density, it can be expected that more cells represent a similar state. At higher values of *k*, we ensure that these are not sampled exhaustively, but rather unique cell states become represented in the random walk.

### Step 2. Construct the transition matrix

Following graph construction, the transition matrix is constructed. We do this by biasing the cell-to-cell transitions to encourage traversal on the graph along a pseudo-temporal axis. This is important to ensure sufficient gene expression patterns are captured across the graph, as these are employed in downstream steps to infer pseudotime. Without bias, random walks are largely limited to traversing the cells surrounding their starting point, and it becomes less likely (enough) meaningful gene expression patterns can be captured. To bias the transitions we require an initial estimate of pseudotime, for which we compute the shortest path distance from a user-defined starting cell population towards all other cells in the data using Dijkstra’s algorithm [20]. Use of shortest-path distance is only suitable when the embedding represents the developmental topology, underscoring the importance of proper dimensionality reduction. Given a starting cell population consisting of *S* cells, a cell’s shortest path distance is computed as:

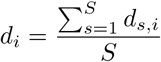

Where *d_s,i_* represents the shortest path distance between cell *i* and starting cell *s*. These distances are computed using the edge weights of the k-NN graph, which represent the distance between cells in the embedded space. This ensures that a shortest-path does not only depend on graph-based shortcuts but is rather also influenced by cell-to-cell similarity. This way, the shortest paths represent probable developmental paths. From computing the shortest path distance, the edge from cell *i* to its neighbor *j* is biased following:

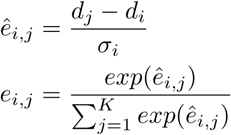

Where *σ_i_* represents the standard deviation over the average shortest path distances of the *k*-nearest neighbors of cell *i*. We apply softmax to further exacerbate the differences between shortest path distances of neighboring cells and thus encourage transitions from cell *i* to cell *j* when *d_i_ < d_j_*. This is needed as, particularly in dense graphs, the shortest-path distance between a cell and its neighbors is small, which is insufficient to bias random walks away from the starting cell. Following biasing, we obtain a directed weighted graph *G* = (*V, E, ϕ*), where *ϕ* represents the weight function defined above.

### Step 3. Perform random walks

Given the transition matrix, we perform random walks on the graph *G*. These random walks are performed to capture gene expression patterns which are represented across the population of cells sampled along the walk. Random walks are initiated separately for each cell in a user-defined starting cell population. We do this not only to increase graph coverage, but furthermore to mitigate bias in the random walks that may be introduced due to their starting position. With a single starting cell, bias that may exist in the paths taken across the graph may cause some cells to not be represented (sufficiently) among the gene expression patterns. In turn, this affects trajectory inference, which is based on these expression patterns. Traversal of the random walk over the graph is governed by the transition matrix described above. Random walks are terminated either when the maximum number of steps, *m*, is reached, or when any cell has been visited more than *t_c_* times during the random walk (see the Supplementary Note for guidance on hyperparameter selection). We set this threshold to prevent unnecessary cycling at the boundaries of the graph and thus save computational time. At *t_c_ ≥* 1, some cycles remain part of the random walk. These are removed by discarding the sequence of cells that intermit the cell that is revisited. We remove these cycles because they represent a cell revisiting a previous, more potent, cellular state, which is considered unlikely. As the transition matrix gets updated (Step 6), transitions that promote cycling should become less probable so that over time progression along pseudotime is enforced. Besides filtering out cycling segments, we further remove random walks shorter than a threshold *t_l_*, to ensure that smooth gene dynamics are captured along a long temporal segment.

### Step 4. Score gene dynamics

Random walks are scored based on the smoothness of the gene dynamics represented by the sequence of cells visited during the walk. Smoothness is selected as a measure because we assume that the expression dynamics that arise from transcriptional regulation in response to differentiation are relatively gradual. Genes that are not actively regulated are expected to either not be expressed, or be expressed noisily, and thus in both cases do not represent a gradual change in expression over pseudotime. By scoring for smoothness, we can differentiate random walks that are likely to represent a developmental trajectory from those that do not. Simultaneously, we identify those genes that may be involved in cell fate specification. As a smoothness measure, the lag-*f* autocorrelation is computed, which measures the Pearson correlation between the gene dynamics and its shifted self. Because a random walk can capture smooth expression patterns for multiple genes, the autocorrelation is computed for all walk-gene combinations. The lag-*f* autocorrelation of walk *w* for gene *g*, *a^f^*, is then defined as:

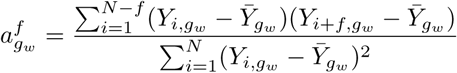

The lag parameter *f* controls the extent to which the gene dynamics is shifted relative to itself. Generally, a slightly higher lag value makes the autocorrelation more robust against random walks where only few (¡ 5) cells have positive expression values.

### Step 5. Align random walks based on smooth gene dynamics

Smoothness by itself is not a sufficient metric to determine the validity of a random walk in terms of capturing a developmental trajectory. To gain more confidence in the biological relevance of the sampled gene dynamics, we compare the random walks to identify the expression patterns that are shared among different walks. Gene dynamics that are captured along multiple, unique trajectories on the graph are assumed to reflect transcriptional dynamics that are involved in the differentiation process. In turn, random walks that capture these gene dynamics are believed to represent possible developmental trajectories. To identify the relevant random walks and expression patterns, the following steps are performed:

- Step 5.1 – Select smooth gene dynamics with *a^f^_gw_ > t_a_*.
- Step 5.2 – Select a seed among the smooth gene dynamics.
- Step 5.3 – Align remaining random walks to the seed and assess alignment for autocorrelation score.

*Step 5.1* Random walks with gene dynamics scoring an autocorrelation above a given threshold *t_a_*_1_ are selected. These random walks are further filtered such that those selected have smooth gene dynamics for at least *t_g_* genes, and each gene has at least *t_r_* random walks with smooth dynamics. We perform these filtering steps to ensure that there are (1) sufficient random walks per gene to compare, and (2) because we assume that multiple genes change dynamically over pseudotime. Random walks that capture multiple smooth dynamics are assumed to reflect a possible trajectory from a starting cell to its end state. A final filtering is applied to ensure that the gene expression is sufficiently dynamic, so that at least *t_d_* cells of the random walks have nonzero expression. This is done to further ensure that the expression pattern arises from active regulation. *Step 5.2* To align random walks for each gene, a representative walk is selected to which the gene expression dynamics of the other random walks are aligned. This is done separately for each gene. Because prior to alignment it is unclear which random walk is most representative of the gene expression pattern represented by the collection of walks that captured smooth dynamics for the respective gene, we consider two criteria to score their quality. First, random walks capturing smooth gene dynamics for multiple genes are preferred, because we assume that these better capture the dynamic changes in gene expression due to differentiation. Second, if multiple random walks exist with an equal number of smooth gene dynamics sampled, the longest random walk is selected as this provides a better baseline to align the other walks to. The selected random walk is hereafter referred to as the seed. In later iterations, it is possible that for a given gene a seed already exists. When this occurs, a candidate seed is selected among the new set of random walks and compared to the existing seed. This is done by attempting to align the candidate to the existing seed(s). If they can be aligned, we assume the candidate does not represent a different gene expression pattern. In this case, a representative walk is selected among the candidate and existing seed(s) based on the aforementioned criteria. If the candidate could not be aligned to the existing seed(s), we assume that it represents a distinct expression pattern. We then select a seed based on the previous criteria and the candidate seed is stored.

*Step 5.3* Having selected the seed, the random walks are aligned. Alignment is performed by computing the autocorrelation over the aggregated dynamics of the seed and the shifted random walk. These shifts are applied as random walks are not always initiated from the same cell and can take slightly different paths over the graph. Comparing the expression patterns without a shift may then return a low autocorrelation score, despite the expression patterns being qualitatively similar. The random walk is shifted maximally *h* positions in both directions relative to the seed. At *h* = 0, the starting cells of both walks are in the same position, while at *h* = *−*1 the starting cell of the aligned random walk is one position to the left of the starting cell of the seed, at *h* = 1 it is one position to the right etc. The shift giving rise to the highest autocorrelation is stored as the best alignment, given that this autocorrelation exceeds the threshold *t_a_*_2_. In doing this alignment, a pseudo-ordering of the cells is created, which is later employed to predict their pseudotime.

Random walks that could not be aligned to the seed at an autocorrelation of at least *t_a_*_2_ can be aligned to a second seed if this exists. If more than *t_w_* random walks remain after alignment, a candidate seed is selected among these random walks. Alignment is then performed only with those random walks that could not be aligned. We perform this additional alignment to improve the ability of the model to capture divergent gene dynamics. Because the seeds are selected partly based on their length, these may be biased to trajectories that are denser. By selecting a seed among those random walks that could not be aligned, we assess whether the population of non-aligned random walks represents a unique expression pattern relative to those aligned. If more than *t_r_* random walks could be aligned to a new seed, these walks and the seed are stored to reflect a unique gene dynamic that can be used for alignment in subsequent iterations.

### Step 6. Update cell-to-cell transitions

To bias the random walks towards the smooth gene dynamics, the transitions visited for the aligned random walks are updated. We do this to encourage traversal along putative developmental paths and thus enhance the ability of the model to capture smooth expression patterns. As zero expression regions do not capture information about the expression dynamics in this region of the graph, only transitions in regions of cells expressing the respective gene are updated. To achieve this, sub-segments of the random walks with zero expression that span at least *t_d_* cells are discarded. The remaining transitions are updated by a value *u*. The magnitude of this value depends linearly on the number of smooth gene dynamics the transition is part of. To ensure that the edge weights still represent probabilities, the weights are rescaled after updating so that the final update magnitude can vary. Across iterations, a transition is only updated once for each unique gene to ensure that the random walks are not biased too strongly towards a subset of gene dynamics which may have a spatial bias. This is especially true in multi-lineage datasets where each lineage is not represented by an equal proportion of cells in the dataset. Updating too strongly could bias the random walks towards the lineage that is most well represented, which would impact trajectory inference as insufficient gene dynamics may get captured for other lineages.

Once the transitions are updated, *Moirai* repeats Step 3 to 6 until a maximum number of iterations, *i*, is reached. By progressively updating the transition matrix based on the aligned gene dynamics, we enhance the ability of the model to capture expression patterns. These aligned gene dynamics collected over multiple iterations are used in the final step of *Moirai*.

### Step 7. Predict pseudotime

In the final step of *Moirai*, the aligned gene expression patterns are aggregated to define a common pseudotime axis along which all cells are ordered. Because cells can be represented across different gene expression patterns, their final pseudotime prediction is an average feature based on their progression along these various gene dynamics. Pseudotime prediction proceeds in six steps (Figure 2):

- Step 7.1 – Retrieve the alignment of the random walks to their respective seeds, and construct the seed-specific alignment matrices.
- Step 7.2 – Infer which random walks were aligned to multiple seeds.
- Step 7.3 – Align the seed-specific matrices using the shared random walks as guides.
- Step 7.4 – For seed-specific matrices that could not be aligned, identify similar random walks and use these for further alignment.
- Step 7.5 – Compute the average position of a cell in the aggregated alignment and rescale to obtain pseudotime.
- Step 7.6 – Predict pseudotime of cells not represented among the aligned random walks.

**Figure 2:**
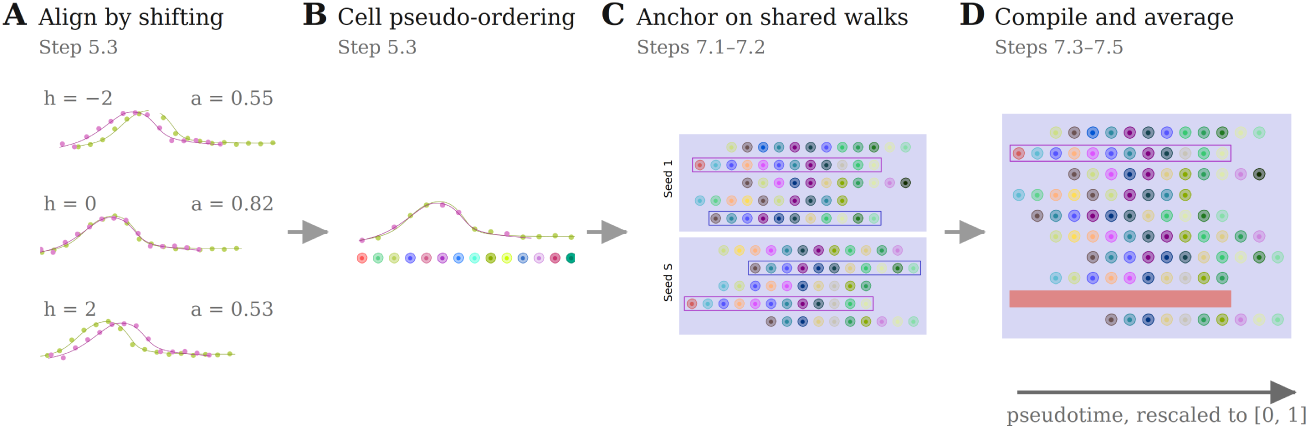
Alignment of random walks and their aggregation into a common pseudotime axis. (A) Random walks that captured smooth dynamics for a given gene are aligned to a representative walk, the seed, by shifting them maximally *h* positions in both directions. The shift giving rise to the highest autocorrelation is stored as the best alignment (here *h* = 0). (B) In doing this alignment, a pseudo-ordering is created of the cell populations sampled by the seed and the aligned random walk. Circles represent cells, colored by their identity. (C) For each seed, an alignment matrix is derived in which every random walk is positioned relative to its seed. Walks aligned to multiple seeds (boxed) are identified, from which a single common walk is randomly selected. (D) Employing these walks as guides, the seed-specific matrices are aligned so that the common random walks take up the same position along the second axis of the final alignment matrix, which thus represents the pseudotime axis. Common walks are removed such that only a single representative is kept, after which pseudotime is predicted as the average position of a cell and rescaled to [0, 1].

*Step 7.1* From aligning the random walks in Step 5 based on the gene expression patterns, we obtained an estimate of the relative positioning of the cells represented in a random walk and its respective seed. To aggregate this information, we derive a L x R x W matrix, *A_s_*, where L is the number of seeds, R the maximum number of random walks aligned to any seed, and W the maximum length of a random walk plus two times the shift (*h*) that was used for alignment. Along the L axis we collect the seed-specific alignment matrices, where every random walk is positioned relative to its seed as determined by the optimal alignment from Step 5. These matrices then each capture a pseudotime estimate for a collection of cells based on the expression pattern represented by the respective seed.

*Step 7.2* To define a single pseudotime axis across expression patterns, we need to compile the seed-specific matrices into a single matrix. To do this, we identify the random walks that are represented in multiple matrices. Because we previously only selected random walks that captured smooth expression patterns for multiple genes, we can expect that some walks were aligned to multiple seeds. This means that for the cells represented in these random walks, we have multiple estimates of their position relative to other cells, as determined for the various expression patterns. By using these random walks as anchors to align the seed-specific alignment matrices, we effectively combine this information across expression patterns. When multiple random walks are shared between two seed-specific alignment matrices, we randomly select a single common walk. Since the overall alignment is performed multiple times, by randomly selecting the set of common walks, we aim to mitigate potential bias of the pseudotime prediction due to the set of walks that were selected to act as guides.

*Step 7.3* Employing the selected random walks as guides, the matrices represented along the L-axis of *A_s_* are aligned. We do this by first identifying the seed-specific matrix that shares the most random walks with other matrices, and setting this matrix as baseline. We then align all other matrices sharing a random walk with this baseline matrix so that the common random walks take up the same position along the second axis of a new matrix *A_f_*, which represents the final alignment matrix. The second axis of *A_f_* will then represent the pseudotime axis, which is fully defined by the gene expression patterns. By updating the set of aligned and non-aligned seed-specific matrices iteratively, we assess if the random walks represented in the newly aligned seed-specific matrices can in turn act as anchor. In this way, we perform this step until no common random walks exist between the aligned and non-aligned seed-specific matrices.

*Step 7.4* For those seeds that could not be aligned using the shared random walks, we search for non-exact matches between the walks of the aligned and non-aligned seed-specific matrices. These non-exact matches represent a continuous subsequence of visited cells in two walks that is at least *t_e_* cells long. As for the alignment based on the common walks, the similar random walks are used to align the seed-specific matrices such that the common subsequences are positioned identically along the second axis of *A_f_*. By iteratively updating the set of aligned and non-aligned matrices, we perform this step until all seed-specific matrices are represented in *A_f_* or no alignment is possible anymore using approximate matching.

*Step 7.5* Once we are done with alignment, we move to pseudotime prediction. To avoid artificially biasing the pseudotime of the cells that were part of the common random walks used for alignment, these walks are removed from the *A_f_* such that only a single representative is kept. Pseudotime is finally predicted as the average position of cell *i* in the final alignment matrix as:

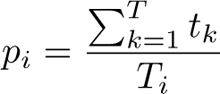

Where cell *i* is represented *T* times in the final alignment matrix. The pseudotime is finally rescaled to [0,1]. To ensure the final pseudotime prediction is not strongly biased by the selection of common random walks used to align the different seeds, we perform all steps multiple times with a different random selection of shared random walks. The pseudotime of a cell is the average over these different runs. To quantify uncertainty in our final pseudotime prediction, we compute the standard deviation over the pseudotime prediction of the different iterations. Furthermore, a mean standard deviation is computed over the positions of a cell in *A_f_* over the different iterations.

*Step 7.6* Cells in areas of the graph that were difficult to reach during random walks or that did not capture smooth gene dynamics are not in the final alignment. To predict pseudotime for these cells, we perform k-NN imputation. Pseudotime is then predicted as the mean pseudotime over the *k* neighbors of a given cell. This is done because we assume that cells that are near in the graph, have a similar developmental state, and so a similar pseudotime.

### Downstream analyses

#### Entropy computation

To assess the developmental potency of cells, we computed the entropy over the probabilities with which cells commit to the different terminal fates. As cells progress along the developmental continuum, they progressively lose their ability to differentiate into multiple fates, eventually becoming restricted to a single lineage. Computing the entropy over the probabilities with which a cell commits to the different terminal states, gives some proxy of its potency. Terminal states are defined manually, through use of marker genes. The probabilities are then computed by performing random walks using the updated transition matrix returned by *Moirai*. As transitions are updated based on the gene expression patterns, we believe that the updated matrix biases walks towards potential developmental trajectories, motivating its use for entropy computation. From each cell in the graph, 1,000 random walks are initialized and terminated when they reach a terminal cell or when the maximum number of steps, *m*, is reached. By counting the number of times cells belonging to each terminal state are visited, we assess the probability with which a cell commits to each terminal fate. Similarly, by computing the entropy over the visit frequencies, we obtain a measure of the potency of the cell.

We performed entropy computation for the myelopoiesis data. Here, we defined four terminal cell states using marker gene expression, which were the plasmacytoid dendritic cells (pDC) (*SCT*, *IRF8*, *RNASE6*, *IGKC*), neutrophil (Neu) (*AZU1*, *ELANE*, *PRTN3*), and erythroid (Ery) (*AHSP*, *APOC1*) lineage. For the fourth lineage, we selected cells expressing the granulocyte-monocyte marker *MPO* and excluded pDC and Neu cells from this selection. We annotated this fourth lineage as granulocyte-monocyte other, as granulocyte-monocyte progenitor cells give rise to both dendritic cells and neutrophils next to other cell types such as macrophages and eosinophils. This fourth lineage was spatially in between the pDC and Neu lineages, and shared some marker gene expression with these lineages, further motivating our annotation.

#### Cluster analysis

To evaluate the gene expression patterns of the genes returned by *Moirai*, we performed cluster analysis. Expression values over the full range of pseudotime were smoothed using moving average, which computes the average value of the expression count over a window of size *b*. The moving average was computed to mitigate the impact of transcriptional noise on the clustering output. Using the smoothed data, Leiden clustering was performed which gave our final clusters representing distinct gene expression patterns [21].

### Datasets

#### Myelopoiesis

The first developmental dataset describes the process of human myelopoiesis and was obtained from the Palantir GitHub. These data capture the 16,106-dimensional transcriptomic profile of 4,142 cells spanning the myelopoiesis developmental continuum. We performed log-normalization and selected the 1,500 highly variable genes before obtaining the embedding using UMAP.

#### Spermatogenesis

Our second dataset captures the process of murine spermatogenesis and was obtained from Weiler and Theis [22]. The data originate from experiments performed by Hermann et al. [23] and profile the expression of 54,144 genes in 1,829 cells sampled from the mice testis. We filtered cells and genes, and performed log-normalization. The 1,500 highly variable genes were selected and the embedding was obtained using UMAP.

#### Zebrafish neural crest development

Our third dataset was obtained from Wang et al. [24] and describes zebrafish neural crest cell (NCC) development. The data capture the expression of 8,012 genes in 697 embryonic cells at six distinct time points during early NCC development. We worked directly with the processed data and ran PCA followed by UMAP using the full transcriptome to ensure sufficient graph connectivity.

#### Acoel embryonic development

Our fourth dataset was obtained from Hulett et al. [25] and describes the embryogenesis of the acoel worm *Hofstenia miamia*. The data profile the expression of 20,935 genes in 8,405 cells of the hatchling juvenile worm. We filtered cells and genes and performed log-normalization. The 1,500 highly variable genes were selected to obtain embeddings using UMAP and diffusion map.

## Results

### The *Moirai* algorithm

Cell differentiation occurs through the organized change in gene expression that effectively pushes a cell along its developmental trajectory. Consequently, the transcriptomic state of a cell marks its progression towards its terminal cellular state. Many trajectory inference methods leverage this signal to predict a pseudotime ordering of cells. Yet, an overemphasis on global transcriptomic changes may impede the model’s ability to resolve early branching points which are captured at finer-scale transcriptomic levels. This may impact the interpretability and understanding we gain from model outputs. Motivated by this, we developed *Moirai*, a graph-based method that first identifies those expression patterns that change dynamically along the developmental continuum and leverages these to define a common axis on which to predict pseudotime. The expression patterns we capture are believed to play some role in cell differentiation and are thus considered informative for the developmental stage of cell. Herein, *Moirai* shifts the focus to genes involved in development when inferring a cell’s progression across the developmental landscape.

*Moirai* processes the scRNA-seq data broadly through six steps (Figure 1). First, the data are embedded in a lower dimensional space. This lower dimensional representation of the data should capture the general developmental topology, to ensure meaningful gene dynamics can be sampled in downstream steps. *Moirai* performs dimensionality reduction using UMAP by default, as it has proven suitable for capturing the global structure of the data [18, 19]. As the performance of UMAP, like other dimensionality reduction approaches, can depend on the data itself [19], *Moirai* supports other embedding methods. From the embedded space, we build a *k*-nearest neighbor (k-NN) graph, where the nodes represent cells and connections are drawn between cells with similar transcriptomic profiles. The graph is then utilized in subsequent steps to perform random walks and sample gene dynamics. To encourage these random walks to move away from their starting point, and up pseudotime, we bias the transitions using shortest-path distance. The transition matrix we obtain through this biasing is used to perform the random walks, which are initiated from a user-defined starting population of cells.

The goal of the random walks is to capture gene expression patterns across the developmental landscape. To assess the quality of these expression patterns, we score the random walks for smoothness. Smoothness is selected as a measure because we assume that genes under transcriptional regulation change more gradually across pseudotime compared to genes that are not expressed or expressed noisily. The scoring is done separately for each gene, so to identify those with smooth expression patterns for a given random walk. *Moirai* then proceeds by comparing the smooth gene expression patterns captured among different random walks. This is done through an alignment, where the expression patterns are shifted and scored for their combined smoothness to assess the most optimal relative positioning. In doing this, the model assesses if an expression pattern is captured by multiple random walks, which increases the confidence that this expression pattern represents a developmental signal.

Through aligning the random walks, we create a pseudo-ordering of two population of cells (i.e., those sampled by the random walks) based on their respective gene expression patterns. By iterating through the various steps multiple times, we assemble a collection of gene-wise pseudo-ordering of cells which we finally aggregate to predict pseudotime. A cell’s pseudotime is then fully set by its average position relative to other cells based on the smooth gene expression patterns.

### *Moirai* captures lineage-specific gene expression patterns of myelopoiesis

Hematopoiesis is the developmental process from which all blood cells derive and which continues throughout an organism’s life. An early fate restriction during hematopoiesis distinguishes the myeloid and lymphoid branches of the hematopoietic lineage [26]. Myeloid cells include erythrocytes and monocytes, while T-, B-, and NK-cells belong to the lymphoid arm. These cells are all widely involved in immune and inflammatory responses, underscoring the importance of their proper development. As hematopoiesis has been well studied, it lends itself to trajectory inference analysis due to the possibility of biological validation. We applied *Moirai* to a developmental dataset spanning the progression from common myeloid progenitors to cells biased towards more differentiated myeloid cell types.

We identified 73 genes for which smooth expression patterns were sampled across the graph. Pseudotime could be predicted leveraging these gene dynamics and aligned well to the general developmental topology (Figure 3A). Due to the use of shortest-path distance as an initial estimate for pseudotime, we were interested to know whether the final pseudotime prediction deviated from this initial guess. We found that final pseudotime prediction was correlated, but not identical to the initial shortest-path distance, illustrating that the model can adjust pseudotime based on the developmental signal represented by the gene dynamics it samples. Comparing *Moirai* pseudotime with that of another graph-based method, Palantir, shows that the two predictions are well correlated despite being predicted from different embeddings (UMAP and diffusion map, respectively) (Figure 3B) [27]. Unlike most pseudotime methods, we further estimated a standard deviation over the pseudotime prediction. Standard deviation was generally low and stable across the pseudotime axis; demonstrating that gene dynamics were sampled relatively uniformly across the graph (Figure S1).

**Figure 3:**
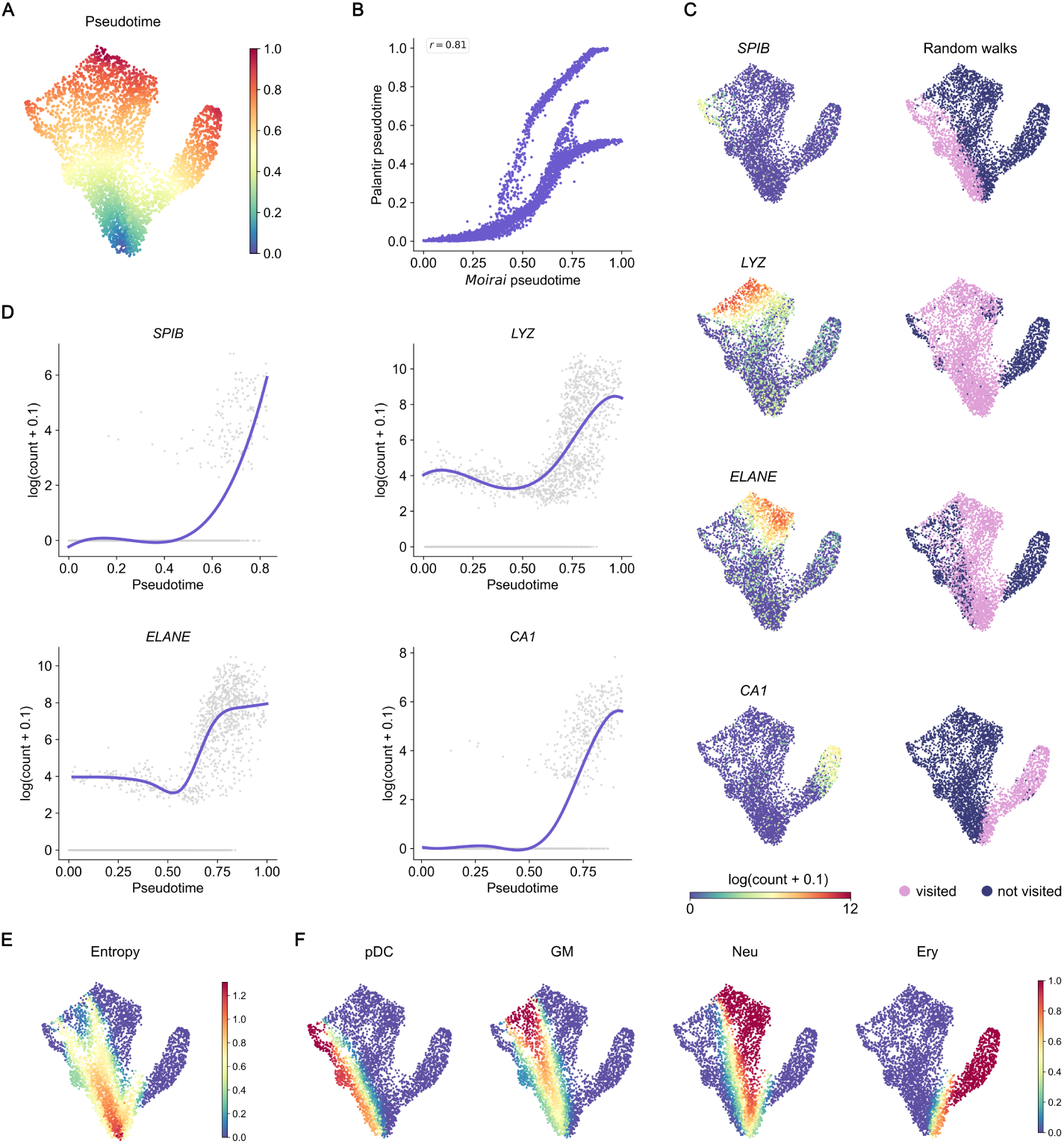
*Moirai* identifies lineage-specific gene expression patterns for four terminal cell states in myelopoiesis. (**A**) UMAP of myelopoiesis dataset colored by *Moirai* predicted pseudotime. (**B**) Correlation of *Moirai*’s and Palantir’s pseudotime prediction. The Pearson correlation is 0.81. (**C**) UMAP of the expression of *Spi-B*, *LYZ*, *ELANE*, and *CA1* together with the coverage of the random walks that captured the expression patterns for the respective genes. (**D**) Expression patterns of the four genes as identified by *Moirai* during trajectory inference. (**E**) Cell entropy computed as a proxy for developmental potency. (**F**) Probability of committing to either of the four terminal fates as computed through performing random walks using the updated matrix returned by *Moirai*. pDC, plasmacytoid dendritic cell; GM, granulocyte-monocyte other; Neu, neutrophil; Ery, erythroid.

Gene expression patterns captured by *Moirai* had clear spatial bias towards specific cell populations, based on which we annotated four global terminal fates: plasmacytoid dendritic cell, neutrophil, granulocyte-monocyte other, and erythroid. Plasmacytoid dendritic cell (pDC) marker genes returned by *Moirai* included *Spi-B*, *IRF8*, and *IRF7* which all have described roles in the development of this subtype of dendritic cells (Figure 3C & D; Figure S2) [28, 29, 30, 31]. Notably, the spatial bias of these genes together with their known role in pDC development suggests a potential branching of the pDCs from the other dendritic fates. Several other genes had clear biasing towards the pDC lineage but had less well-described roles in this developmental process. *C12ORF75* was lowly expressed in cells across the developmental continuum but appeared upregulated in cells biased towards the pDC lineage (Figure S2). This gene has been studied as a marker of cancer progression where it was found to positively correlate with the migration of immune cells, including dendritic cells [32]. These findings may suggest some role of *C12ORF75* in pDC migration. Its expression coincides with that of *SCT* with a role in the induction of immature dendritic cell chemotaxis, further suggesting the development of cell motility (Figure S2) [33].

*ELANE* was among the genes returned for the neutrophil lineage and has been studied for its role in neutrophil maturation (Figure 3C & D) [34, 35]. Alongside *ELANE*, smooth expression patterns were returned for *AZU1*, *PRTN3*, and *CFD* which are clustered together spatially on chromosome 19 (Figure S2) [36]. The former three genes are all primary components of neutrophil azurophilic granules, while *CFD* is secreted. Comparing the distribution of these four co-regulated genes with that of the granulocyte-monocyte marker *MPO* [37], highlights a potential branching of the neutrophilic lineage from the other granulocyte-monocyte fates (Figure S2). This branching is further highlighted by *LYZ*, whose expression marks the cells sitting between the neutrophil- and plasmacytoid dendritic-like cells (Figure 3C & D). *LYZ* encodes a glycoside hydrolase, and in zebrafish was found specifically expressed by macrophages, which derive from the granulocyte-monocyte progenitor and are closely related to dendritic cells [38, 39]. The gene expression patterns are well captured by the random walks, illustrating the ability of *Moirai* to capture relevant gene expression patterns specifically and completely. Notably, *Moirai* is able to do so even when the bifurcation is not captured in the graph topology. A likely consequence of this was a higher standard deviation of cells belonging to those trajectories that captured gene expression patterns for the neutrophil and granulocyte-monocyte other lineage (Figure S1). This highlights the increased ambiguity of cell fate commitment near a branching point.

We finally recovered several genes specific to the erythroid lineage, including *CA1* which has been linked to erythroid differentiation (Figure 3C & D) [40]. *CA1*-deficient cells had impaired expression of globin-related genes, suggesting some involvement of *CA1* in heme and hemoglobin synthesis. We similarly identified *AHSP* with an expression pattern limited to the erythroid lineage (Figure S2). This gene has been marked as an erythroid-specific gene where it acts as an *α*-globin-specific chaperone [41]. *BLVRB* was found with an expression pattern similar to that of *AHSP* and has been linked to erythrocyte and megakaryocyte lineage commitment (Figure S2). *Moirai* additionally identified *APOC1* with erythroid-specific expression (Figure S2). This gene has been implicated in the M2 polarization of macrophages and was reported as a biomarker of cancer progression [42]. *APOC1* is believed to be involved in lipoprotein metabolism, but its exact function remains unclear. Lipoprotein metabolism has been marked as important for erythroid cell maturation and a related gene, *APOA1*, has already been identified as a terminal erythroid marker [43, 44].

From the gene dynamics returned by *Moirai*, we finally estimated the developmental progression and thus bias towards each of the lineages. By updating the transitions part of smooth gene dynamics, we assume that paths become biased towards meaningful developmental trajectories. Through initiating random walks from all cells in the updated graph, we simulate their probable developmental paths and obtain an estimate of the probabilities with which they commit to each terminal state. Computing the entropy over these probabilities, we get a proxy for the potency of the cell, which captures its developmental potential. We found entropy to be highest for cells with low pseudotime and to decrease gradually towards the terminal cell states (Figure 3E). The earliest drop in entropy belonged to the erythrocytic lineage, which is in line with its developmental branching from the granulocyte-monocyte lineage. Around mid pseudotime, cells already showed clear bias towards either of the three remaining fates, despite this branching not being captured in the graph representation (Figure 3F).

### Cluster analysis reveals windows of transient upregulation of genes involved in spermatogenesis

Underlying male fertility is the developmental process of spermatogenesis, during which germ cells differentiate to form mature reproductive cells. Mouse spermatogenesis takes place in the seminiferous tubules, where cells across the developmental continuum reside [45, 46]. The developmental process begins with the differentiation of germ cells into type *A* spermatogonia. The spermatogonia go through mitosis to differentiate from type *A*_1_ to type *A*_4_ spermatogonia. Some *A*_4_ spermatogonia further differentiate to form type *B* spermatogonia which give rise to the primary preleptotene spermatocytes. In order, these further differentiate to form the leptotene, zygotene, pachytene, and diplotene primary spermatocytes. Through two rounds of meiosis, the diplotenes go on to form the round spermatids which then undergo a transformation during which the acrosome (Golgi-derived structure found in the head of spermatozoa) and flagellum are formed. These elongating spermatids finally produce the spermatozoa, which are released into the lumen. The developmental continuum of cells residing in the mouse testis, marks it as a suitable system to study for trajectory inference. With *Moirai*, we studied a developmental dataset capturing the expression of 1,829 cells ranging from spermatogonia to round spermatids [23]. We captured a total of 371 genes with smooth expression patterns across the developmental landscape and inferred pseudotime employing these (Figure 4A). Pseudotime prediction aligned well with the known progression of developmental stages and correlated strongly with the shortest-path distance (Figure 4B; Figure S3A). Such correlation may be expected in datasets with a nonbranching developmental landscape, where cells are likely to be only nudged slightly through the alignment of gene expression patterns. Despite all cells being visited by the random walks (Figure S3B), pseudotime could be predicted for only 93% of cells. The population of cells for which no pseudotime was predicted showed a distinct transcriptomic state compared to other round spermatids, with upregulation of genes found highly expressed in spermatogonia (Figure S4). The transcriptomic signature may mark this population as separate from the remaining round spermatids, which could explain the lack of gene expression patterns captured for cells belonging to this population.

**Figure 4:**
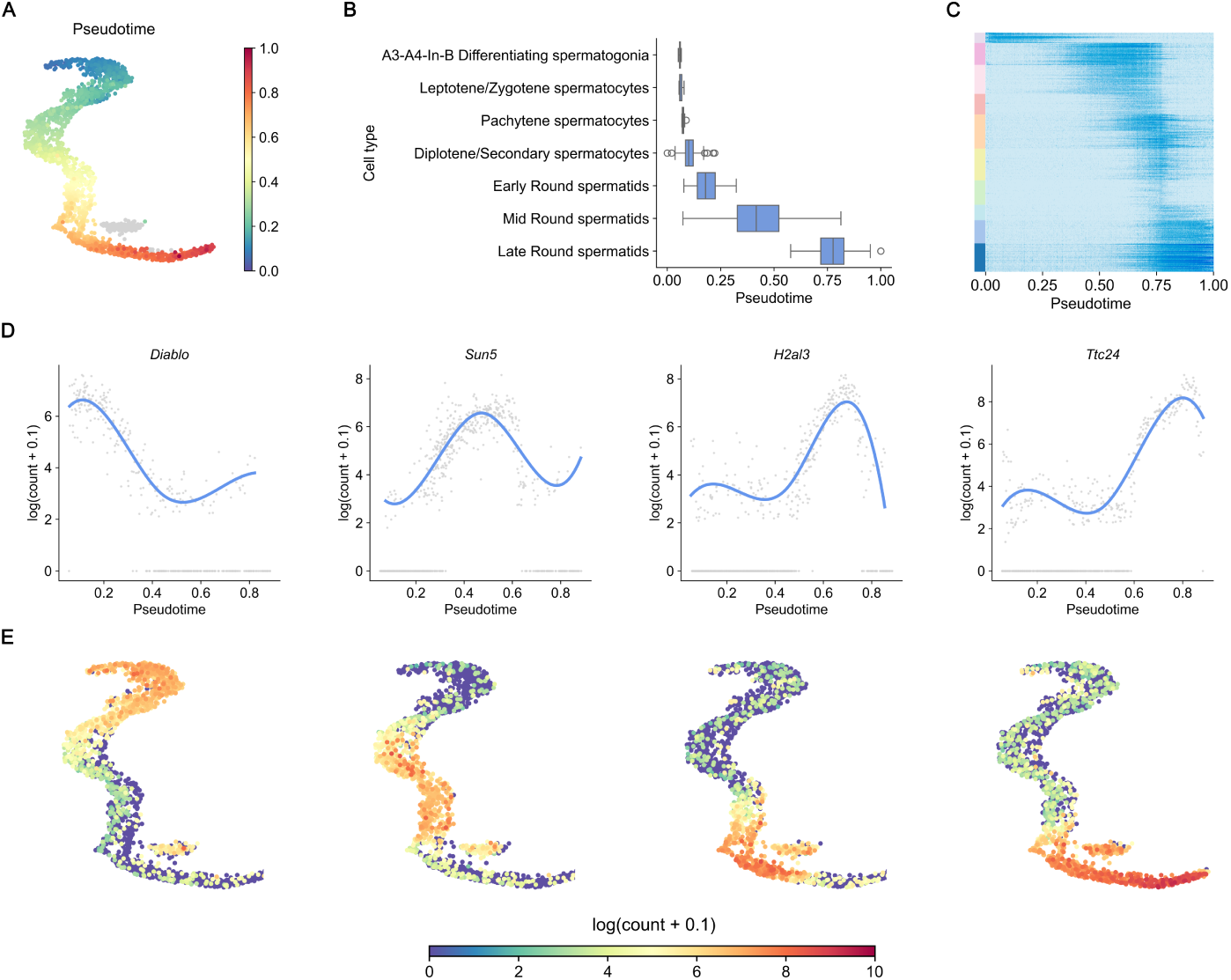
Gene expression patterns captured for spermatogenesis display transient windows of upregulation during round spermatid pseudotime. (**A**) UMAP of spermatogenesis dataset colored by *Moirai* predicted pseudotime. (**B**) Boxplot of *Moirai*’s pseudotime and the developmental stages of spermatogenesis. (**C**) Heatmap showing gene expression of 371 genes returned by *Moirai* over pseudotime. Bar on the left of the plot indicates the ten clusters identified with cluster analysis. (**D**) Expression patterns of *Diablo*, *Sun5*, *H2al3*, and *Ttc24*, which belong to clusters with distinct regions of expression across pseudotime. (**E**) UMAP showing gene expression of the four respective genes.

To evaluate the gene expression patterns captured by *Moirai*, we performed a cluster analysis. Gene expression patterns could be assigned to ten unique clusters with distinct windows of expression across pseudotime (Figure 4C; Figure S5). Genes downregulated early in pseudotime were mainly involved in cell cycle regulation, with cluster representatives including *Aurka* and *Nek2*, which both have been found involved in meiosis I (Figure S6) [47, 48]. We additionally found the pro-apoptotic gene *Diablo* to be highly expressed in spermatogonia, and its expression appeared to be slightly upregulated again later in pseudotime (Figure 4D & E). *Diablo* is involved in the mitochondrial apoptotic pathway and has previously been found expressed in spermatocytes and spermatids [49]. Apoptosis is central to proper sperm cell development [50], and we annotated further genes relating to this process across pseudotime.

Most genes were expressed only transiently, coinciding with distinct phases of round spermatid pseudotime. *Sun5* was expressed in a window spanning the progression from early- to mid-phase round spermatids and is essential for proper development of sperm morphology (Figure 4D & E) [51]. Around this time frame we additionally observed upregulation of *Catsper3* and *Prr27*, which are required for sperm motility and acrosome formation, respectively (Figure S6) [52, 53]. Expression of *H2al3* covers the transition from mid- to late-phase round spermatid pseudotime (Figure 4D & E). This testis-specific histone variant may mark the chromatin reorganization that occurs with spermatid maturation [54]. *H2al3* expression coincides with that of a second H2 variant, *H2al2a* further illustrating this (Figure S6). We additionally identified two gene clusters that were expressed during a particularly narrow window in late spermatid pseudotime (Figure S5). Among these was *Cyp2a12*, a gene involved in bile acid production that was identified similarly in the original publication (Figure S6) [23, 55].

We finally recovered two clusters of genes that were upregulated towards the end of pseudotime, marking the maturation of the late-phase round spermatids. Among these genes, we identified *Ttc24* which is a commonly used marker for elongating spermatids, highlighting the transition from round to elongating spermatids towards the end of pseudotime (Figure 4D & E) [56]. Several other genes were related to sperm morphology, such as *Spatc1* which stabilizes the sperm head-tail junction, and *Tex46* which was found essential for sperm head formation and thus mouse fertility (Figure S6) [57, 58]. We additionally identified some genes involved in the oxidative stress response to be upregulated towards the end of round spermatid pseudotime. These genes were *Osgin1* and *Glul*. Though the former gene has not been explicitly linked to sperm cell development, it has been linked to oxidative stress regulation and autophagy in the human airway epithelium (Figure S6) [59]. *Glul* was found to be essential for a proper response to oxidative stress during spermatogenesis [60]. Spermatozoa have been found to activate autophagy when exposed to oxidative stress, and proper regulation of this stress response is required to maintain sperm cell quality [61].

Notably, among the 371 genes returned by *Moirai* we found various genes with an undescribed role in spermatogenesis. Some of these genes encoded putative long non-coding RNAs (lncRNAs), which have been previously implicated in various functions in spermatogenesis across developmental pseudotime [62]. Expression of putative lncRNAs was found across the developmental continuum, with no clear temporal clustering. *Malat1* encodes an lncRNA involved in cell-cycle regulation and was expressed transiently by mid-stage round spermatids (Figure S6) [63].

### *Moirai* resolves pseudotime for non-continuous developmental data

Neural crest cells (NCCs) are transient, migratory cells originating from the dorsal neural tube [64]. Differentiating NCCs can give rise to various cell types including melanocytes, neurons, and smooth muscle cells. The developmental fate of NCCs depends on their location along the main axial level of the embryo, and distinct transcriptional programs regulate the respective cell fates. Many insights into NCC development have been derived from studies on zebrafish, where the possibility of imaging together with genetics has enriched the potential for discovery [65]. We applied *Moirai* to an NCC developmental dataset that captures the transcriptomic profile of 697 cells from the zebrafish embryo [24]. Cells were sampled across six time points spanning the early phases of NCC development. The number of cells captured in this dataset together with their temporal separation due to sampling time points, mark this dataset suitable to assess *Moirai*’s performance when its key assumption of developmental continuity is challenged.

*Moirai* recovered a total of 166 gene dynamics across the developmental continuum. Pseudotime could be predicted for all cells and was largely in line with the sampling time point (Figure 5A). Though this can be partly attributed to the accuracy with which the embedding captured the developmental progression, the final pseudotime prediction was not identical to the initial shortest path distance (Figure S7A). Again, this highlights that *Moirai* is able to leverage gene expression patterns to infer a cell’s position along a pseudotime axis. The standard deviation over the pseudotime shows some spatial bias, with cells of the neural plate border, pharyngeal arches, and the facial mesenchyme lineage having a relatively higher standard deviation (Figure S8). This effect is partly attributable to the lower frequency with which gene expression patterns were captured for these lineages. When more gene expression patterns are captured for a given cell, the confidence in that cell’s positioning may increase, which would be reflected in a lower standard deviation. Indeed, cells that are visited more as part of the gene expression patterns tend to have lower standard deviations (Figure S7B & C).

**Figure 5:**
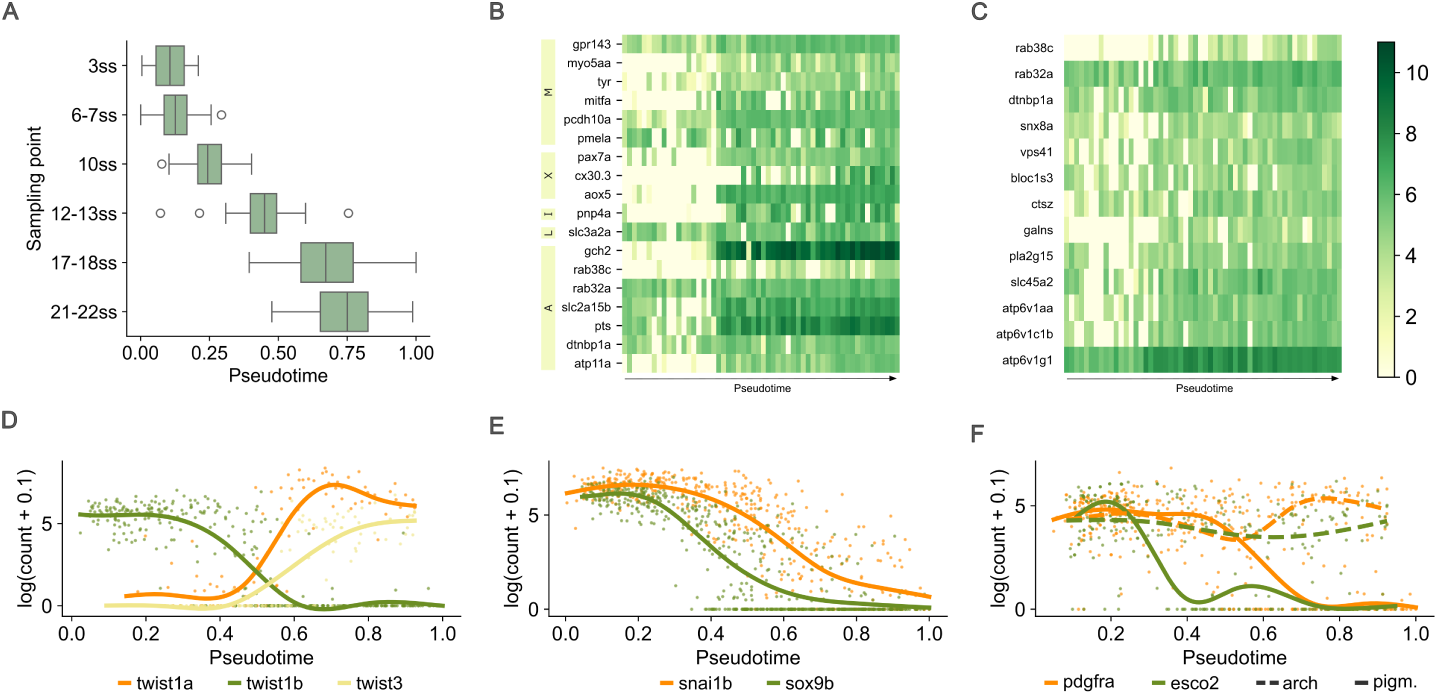
*Moirai* pseudotime aligns well with sampling time of zebrafish neural crest cells. (**A**) Boxplot of *Moirai*’s pseudotime and the sampling time of zebrafish neural crest cells. (**B**) Heatmap of expression (lognormalized) of genes involved in chromatophore development, labeled by chromatophore subtype. (**C**) Heatmap of expression (log-normalized) of genes related to development of lysosome-related organelles. Note that the colorbar is shared between B and C. (**D**) Expression patterns of three *twist* genes captured by *Moirai*. Though the dynamics of *twist1a* and *twist3* were captured from the progenitor towards the head mesenchymal cells, for *twist1b* this was towards the pigment cells. (**E**) Downregulation of *snai1b* and *sox9b* across zebrafish neural crest pseudotime. (**F**) Unique gene dynamics for *pdgfra* and *esco2* for the trajectory towards the pigment cells (solid line) versus the pharyngeal arches (dashed line). Ss, somite stage; M, melanophores; X xanthophores; I, iridophores; L, leucophores; A, ambiguous (i.e., genes found expressed by multiple chromatophore subtypes).

In total, *Moirai* returned 75 expression patterns specific to the pigment lineage. Several genes had described roles in specific subtypes of chromatophores such as melanophores (*gpr143*, *myo5aa*, *tyr*, *mitfa*), xanthophores (*pax7a*, *cx30.3*), leucophores (*slc3a2a*), and iridophores (*pnp4a*). Other genes were more ambiguously linked to chromatophore subtypes. For instance, *gch2* and *slc2a15b* have been linked to both xanthophore and leucophore development [66, 67]. Expression of these genes across the pigment cells was widespread, and no clear subpopulations could be distinguished (Figure 5B). Instead, the clearest separation in the population was due to pseudotime, with some genes being induced later. Such findings either suggest a more general role of these genes in pigment cell development or highlight some ambiguity in expression of cell type-specific genes prior to fate commitment. Indeed, experimental evidence suggests that *mitfa*-expressing cells can still commit to an iridophore fate, despite *mitfa*’s central role in melanophore development [68]. These findings highlighted that cell plasticity is not necessarily reduced by the expression of cell-type specific markers.

Several genes with expression specific to the pigment lineage had undescribed roles in the functioning of these cells. For some of these genes, we hypothesize a putative role in lysosome development. Lysosome-related organelles (LROs) are a diverse group of cellular compartments which find their origin in the endosomal-lysosomal system [69]. Melanophores, xanthophores, and iridophores all contain LROs, which are called melanosomes, xanthosomes, and iridosomes, respectively. Besides genes with known involvement in the development of these LROs (*rab38c*, *rab32a*, *slc45a2*, *dtnbp1a*), we found some with a putative role in LRO development that have not been clearly linked to pigmentation. Three genes *atp6v1c1b*, *atp6v1aa*, and *atp6v1g1* encode subunits of V-type ATPases, and were co-expressed with known LRO markers (Figure 5C). V-type ATPases are involved in maintaining pH levels in melanosomes and iridosomes, which is critical to pigmentation [70, 71]. Although co-expression is not definite evidence of the involvement of these genes in the formation of LROs, it does warrant further investigation.

A second subset of genes was identified with expression limited to cells of the pharyngeal arches and facial mesenchymal cells. The pharyngeal arches are formed from cranial neural crest cells and are numbered according to their spatial positioning along the anterior-posterior axis [72]. The first two pharyngeal arches form the jaw and its support structures. Mesenchymal cells originating from the neural crest populate the pharyngeal arches. Random walks captured expression patterns across these two cell types. Expression of *itga8* was found in cells belonging to the first pharyngeal arch and facial mesenchyme cells (Figure S9). Reports have shown *itga8* expression at the pharyngeal arches, where it was hypothesized to be required for epithelial-mesenchymal interaction that in turn sculpts the facial shape [73]. We also recovered gene expression patterns for three out of four *twist* genes, which are involved in formation of the craniofacial skeleton [74]. Although expression of *twist1a* and *twist3* was more limited to the arch mesenchymal cells, *twist1b* was additionally highly expressed in the progenitor cells while being selectively repressed in the pigment cells (Figure 5D).

A final subset of genes was expressed most strongly in the progenitor cells and was downregulated along the progression of pseudotime. Broader expression of these genes across the developmental continuum highlighted that despite the temporal separation of the cell populations, gradual changes could still be captured. Among this final subset of genes was *sox9b*, a marker of neural crest precursors that becomes downregulated when neural crest cells begin to migrate [75]. We similarly identified *snai1b*, with a slight delay in its downregulation compared to *sox9b* (Figure 5E). The *snai1b* gene is required for epithelial-to-mesenchymal transition of neural crest cells and thus for these cells to acquire their migratory properties [76]. Other genes were only repressed in particular lineages, such as *pdgfra* and *esco2* which have been linked to craniofacial development and cell cycle regulation, respectively, and were specifically repressed in pigment cells (Figure 5F) [77, 78].

### Gene dynamics are robustly recovered across different embedding methods

With different dimensionality reduction techniques, we can expect to generate qualitatively different embeddings for the same data. Due to the dependence of *Moirai* on the embedding for its final pseudotime prediction, we wanted to understand how *Moirai’s* output would change depending on the dimensionality reduction method employed. We did this by embedding developmental data in a lower dimensional space using either UMAP or diffusion map, before running *Moirai* to infer pseudotime. The developmental data used described the embryogenesis of the acoel worm *Hofstenia miamia*. Acoels are marine worms, which are often studied for their ability to regenerate. The dataset covers the transcriptomic profile of 8,402 cells of hatchling juvenile acoel worms, ranging from pluripotent neoblasts to more differentiated cell types [25].

*Moirai* returned 379 genes for the UMAP versus 348 genes for the diffusion map, with most genes shared between the two outputs (Figure 6A). Comparing the random walks recovered for each gene expression pattern showed a large overlap between the two embeddings in terms of cells visited (Figure 6B). In general, random walks from the diffusion map showed a slightly lower overlap, indicating that these walks typically captured additional cells relative to those of the UMAP. When assessing the cell coverage of the gene expression patterns for which the overlap between the random walks was below 50%, it becomes clear that limited overlap only arises due to a higher coverage in one embedding (e.g., as for *s28a1*, Figure S10A) or the expression pattern being captured over a longer range (e.g., as for *wdr52-2*, Figure S10B). Due to *Moirai* aggregating information across multiple expression patterns, full coverage of cells expressing the respective gene is not necessary, provided that sufficient gene dynamics are captured in each region of the graph. As an effect, these inconsistencies should not severely impact the final pseudotime predictions.

**Figure 6:**
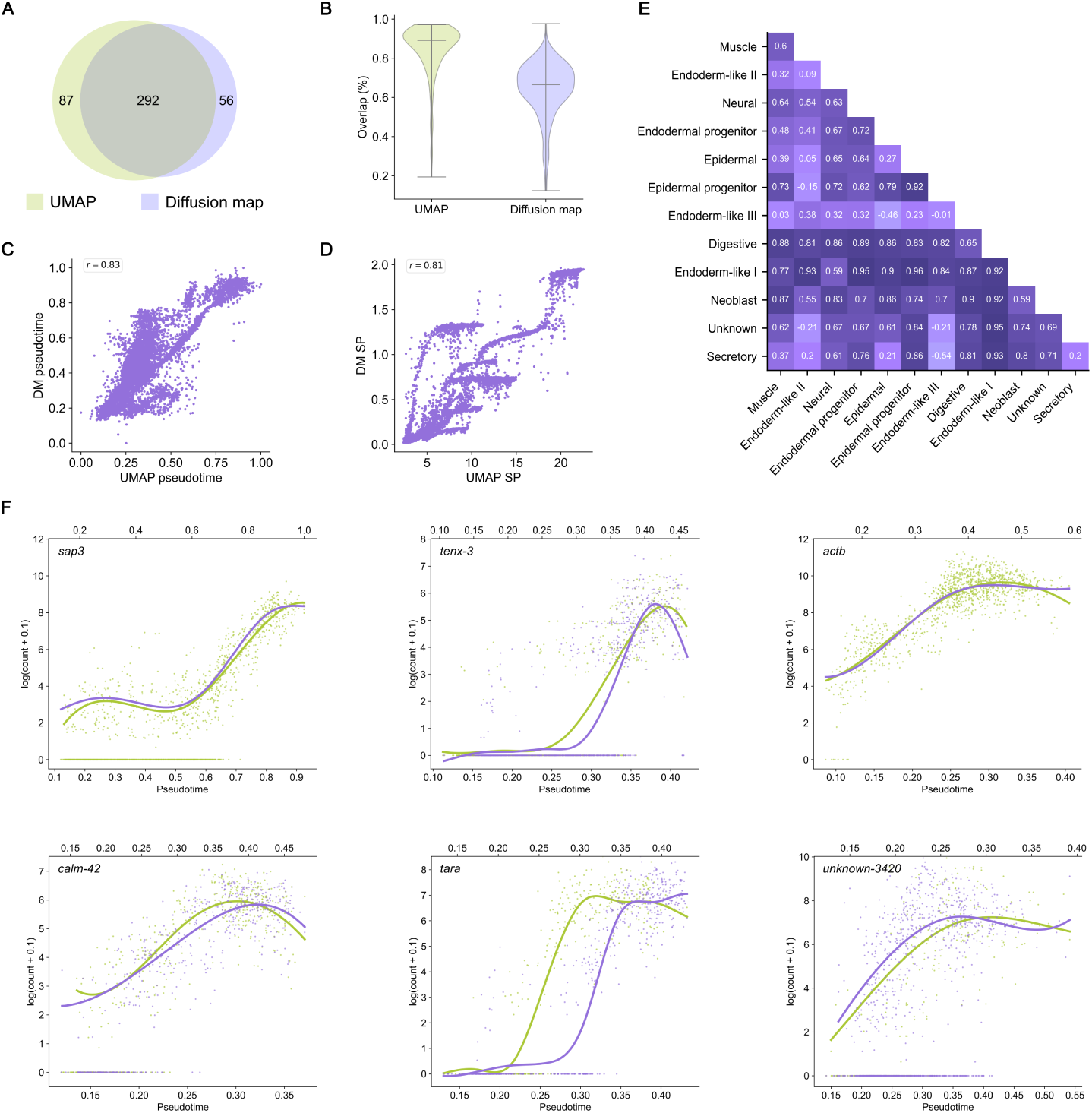
*Moirai* pseudotime changes depending on initial embedding, but expression patterns are consistent. (**A**) Venn diagram of genes returned by *Moirai* when using UMAP versus diffusion map as embedding technique. (**B**) Violin plot of the overlap in cells captured per gene expression pattern for both embeddings. (**C**) Correlation plot of *Moirai*’s pseudotime for UMAP versus diffusion map derived predictions. (**D**) Correlation plot of initial shortest-path distance for both embeddings. (**E**) Spearman correlation of the pseudotime of both embeddings for different pairs of cell types. (**F**) Expression patterns of *sap3*, *tenx-3*, *actb*, *calm-42*, *tara*, and *unknown-3420* captured by *Moirai* for both embeddings. Bottom x-axis indicates pseudotime axis for the UMAP, while the top x-axis indicates that for the diffusion map. DM, diffusion map; SP, shortest path distance.

Pseudotime prediction arising from the gene expression patterns is similar between the two outputs, though not perfectly correlated (Figure 6C). Deviations in pseudotime prediction are most notable on a cell-type level, with some cell types displaying lower correlation between the two outputs (e.g., secretory, endoderm-like II / III) compared to others (e.g., epidermal progenitor, endoderm-like I) (Figure S11). Interestingly, correlation between the initial shortest-path distances of the two embeddings was slightly lower than that of the final pseudotime prediction (*ρ* = 0.81 vs *ρ* = 0.83, respectively), demonstrating that employing the gene expression patterns, *Moirai* was able to define a common axis of pseudotime (Figure 6D). For both embeddings final pseudotime prediction deviated more strongly from initial shortest-path distance than for prior datasets, further demonstrating *Moirai*’s ability to leverage the developmental signal represented by the expression patterns to inform pseudotime prediction (Figure S12).

Because pseudotime is most meaningful between a cell and its progeny, we additionally computed the Spearman correlation between the two outputs for different pairs of cell types (Figure 6E). This should illustrate whether the relative ordering is preserved between the two outputs, and which pairs are most different. The correlation score for pairs involving neoblast were generally high, with lower values being partly attributable to lower Spearman correlation of the second cell type. For instance, Spearman correlation of neoblast-endoderm like II cells was the lowest at 0.55, with the Spearman correlation of just the endoderm-like II cells sitting at 0.09. Other pairs, such as epidermal progenitor-epidermal, showed relatively high Spearman correlation, motivating relatively consistent positioning of these cells along the pseudotime axis. Digestive cells similarly showed relatively high Spearman correlation across the different pairs. To gain a more detailed view on the outputs of each embedding, we finally assessed the expression patterns. Despite the pseudotime differing slightly in magnitude, expression patterns are relatively similar between the two outputs. We found *sap3* similarly upregulated in both outputs, with only a slight deviation in the pseudotime axis (Figure 6F). Notably, in both outputs we identified a subset of *sap* genes upregulated in the digestive cells, suggesting some role for this gene family in the digestive tract. For the neural cells expression of *tenx-2* similarly overlapped, and additional identification of *tenx-3* upregulated in this cell population points to a role for these genes in neural fate specification (Figure 6F; Figure S13). Strong upregulation of *actb*, encoding *β*-actin, was also consistently identified between the two outputs. For the epidermal lineage, we identified *calm-42* with similar expression patterns across embeddings (Figure 6F).

For the secretory lineage, fewer genes were identified, and thus in common, between the two outputs. Expression patterns were generally similar, as displayed by *tara*, though slightly more noisy compared to the other lineages (Figure 6F). Gene dynamics were most different for the endoderm-like II / III lineages. Though expression patterns of *unknown-3420* were relatively similar, most genes showed noisy, dissimilar expression patterns. The positioning of these cell types in the different embeddings was different, with the UMAP capturing more of the bifurcation, while the diffusion map embedded these two cell populations more sequentially (Figure S14 & S15). In addition, the diffusion map further connected these endoderm-like cells to digestive cells. As a result, for genes with expression limited to the endoderm-like II / III cells, diffusion maps tended to capture transient patterns, while UMAP would show only an upregulation. For both outputs, fewer gene dynamics were captured in this region of the graph, which may further suggest that the developmental topology was poorly captured in the embeddings. In turn, this may also have affected the pseudotime prediction and explain why a lower correlation was found for this cell type.

## Discussion

In this report, we presented *Moirai*, a graph-based trajectory inference method that leverages gene expression patterns to define a common pseudotime axis between all cells. *Moirai* was applied to four developmental datasets, where we demonstrated *Moirai*’s ability to predict pseudotime and infer those expression patterns that change dynamically over the graph specifically, and completely. In doing so, *Moirai* shifts the focus to those genes with a likely involvement in cell fate specification and lets these guide a cell’s pseudotime prediction. Our findings validated the involvement of known actors, while identifying other genes with lineage-specific expression with no prior link to the respective developmental processes. We furthermore demonstrated the suitability of *Moirai*’s output for a range of downstream analyses, such as entropy computation and cluster analysis, which further enriched biological interpretations.

The identification and use of gene expression patterns during the inference process sets *Moirai* apart from most pseudotime inference algorithms, which instead employ global transcriptomic changes and mathematical methods [79]. Although *Moirai*, like these methods, summarizes the global topology of the data through dimensionality reduction and graph construction, it then effectively zooms in to individual gene dynamics and how these are formed across the graph. *Moirai* then aggregates these gene dynamics to define a common pseudotime axis across all cells. This way, these dynamics are not only a source of validation through downstream analyses but also guide the process of trajectory inference directly. *Moirai* thus effectively incorporates these small scale biological signals into the inference process. In addition, through aggregating information across different gene expression patterns we gain a measure of uncertainty which is not just linked to the stochasticity of the method itself but is also informed by a cell’s developmental progression as determined by the expression patterns.

Another key feature of *Moirai* is the use of shifts to compare gene expression patterns. The benefit of this feature is twofold. First, it makes the method more robust to the choice of starting cell. To capture gene expression patterns, random walks are initiated from a starting cell population. However, it is unlikely that all cells in this population represent the same cell state. Based on the expression patterns sampled by the random walks that were initiated from each starting cell, *Moirai* infers the relative positioning of these starting cells. As an effect, *Moirai* is relatively robust against the selection of starting cells, which is especially valuable for datasets where selection of the starting cell population is difficult. The second benefit we derive from the shift is that it allows *Moirai* to detect distinct expression patterns for the same gene. These expression patterns may be similar but temporally separated or are dissimilar in their form (e.g., transiently upregulated, versus upregulated). When two gene expression patterns cannot be aligned, these are saved as two separate seeds which are returned by *Moirai*. This way, *Moirai* identifies divergent gene expression patterns during the inference process, instead of only deriving these from downstream analyses. Simultaneously, through assessing the random walk coverage for a given gene expression pattern, *Moirai* similarly identifies gene expression patterns that are shared across distinct lineages. The ability of *Moirai* to directly infer such (divergent) dynamics further sets it apart from other methods.

To further enhance *Moirai*’s ability to recover relevant developmental trajectories, *Moirai* can be extended by incorporating domain knowledge into the scoring procedure. Random walks are scored on a gene-specific basis, with only a threshold to ensure multiple gene expression patterns are captured per random walk. However, we do not consider which gene expression patterns are captured per random walk. Recognizing transcription factor–target pairs, or genes that are involved in the same pathway, may further enhance our scoring system and thus *Moirai*’s ability to discern biologically meaningful trajectories. Simultaneously, it may enhance the model’s ability to gain mechanistic insights into the developmental processes. Most pathway analysis implementations cluster the data, which is useful to coarsely infer which biological processes are active, but is limited in its ability to assess how these processes organize themselves during dynamic processes such as differentiation. Incorporating pseudotime in the inference process, may grant a more detailed view on which pathways are involved in lineage specification, and how these organize themselves sequentially along a pseudotime axis. In addition, assessing the content of a random walk as a whole may also better allow the detection of convergent trajectories which are characterized by a distinct profile of gene expression patterns across pseudotime. Another extension to *Moirai*’s architecture may be imagined for the downstream analyses. We performed various downstream analyses, demonstrating the suitability of *Moirai*’s output for biological validation and interpretation. Expanding on *Moirai*’s architecture may furthermore allow the direct identification of lineages based on the spatial bias in the gene expression patterns. Across three developmental datasets, we found lineage-specific gene expression patterns which were captured by random walks with clear spatial bias. In the current installation, manual annotation of terminal states was required to compute entropy and bias towards these states and gain some indication of lineage commitment. However, leveraging the spatial overlap between subsets of gene expression patterns may allow the model to directly identify putative lineages and which genes are most involved in their specification. This would be especially valuable for systems that have been less explored, and so potential for validation and manual annotation of terminal states becomes limited. Still, it should be noted that though the expression patterns identified by *Moirai* may distinguish lineages, these branching points may not be taken as true bifurcation points. Experimental evidence suggests that cells commit to their respective lineage before this clearly manifests in the transcriptional profile [80]. This could involve a role for epigenetic regulation, a dimension that may further enrich pseudotime inference methods.

Despite the aforementioned strengths of *Moirai*, limitations remain. A main limitation pertains to the approach through which the transition matrix is obtained. Utilizing the shortest-path distances to bias the transitions can be done in various ways, the current implementation thus being a heuristic. Without sufficient biasing of the transition matrix, random walks may be limited to the area surrounding the starting node. Due to the dependency of *Moirai* on the gene expression patterns being properly sampled across the graph, efficient traversal of the graph is fundamental to its performance. As a result, when the graph is biased less severely, many more random walks are required to ensure sufficient expression patterns get captured. Because existing bias in the graph is not expected to impact final pseudotime prediction strongly, the severity of this shortcoming depends on the intended use of *Moirai*. For instance, lineage assignment will be influenced by existing bias in the graph, which may not be necessarily meaningful. For future applications, it may then be interesting to consider other methods of biasing the transition matrix and the impact this has on downstream analyses such as lineage assignment.

Another limitation of the method relates to the contingency of the final pseudotime prediction on the dimensionality reduction method. Running *Moirai* with embeddings generated using UMAP and diffusion map, we found that despite great overlap between the gene expression patterns, pseudotime was not perfectly correlated. Though for some cell types, this was attributable to the qualitative placement of the cells relative to other cell types (i.e., endoderm-like II / III), for others quantitative differences in final pseudotime prediction were related more directly to the embedding approach. While UMAP captures the global topology of the data, in a diffusion map the data embedding is derived from the scaled eigenvectors of the transition matrix [18, 81]. As a result, UMAP representations tend to be less sparse than that of diffusion maps, where cells are represented along narrow lines in a multi-dimensional space. Effectively, this may have resulted in fewer steps being required in the UMAP compared to the diffusion map to reach terminal states of some lineages. Due to the direct relation between the length of a random walk and the final pseudotime prediction, this may have resulted in the discrepancy observed. Without critically assessing the data, this could lead to false conclusions regarding the progression of cells along the developmental path. For instance, we found digestive cells were assigned the highest pseudotime, which may suggest that these cells had progressed further in pseudotime compared to those of other lineages. However, we should consider how meaningful cross-lineage comparisons are, as these are governed by distinct transcriptional programs.

This broadens the discussion to the general goal of pseudotime prediction and the conclusions we may draw from the results. Not only the embedding approach, but other pre-processing choices (e.g., (log-)normalization, cell-cycle correction), impact the embedding we obtain, the results we get, and so perhaps even the conclusions we draw [9]. Though deriving from a common origin, at some point progression of cells is dictated by factors unique to their fate. When developmental progression is governed by different factors, quantitative comparisons of pseudotime may become meaningless. The value that we derive from pseudotime relates to our ability to discern lineages and infer which gene expression patterns are involved in cell fate specification. Pseudotime is then most relevant in how it is defined between cells biased towards the same fate, rather than those with unique destinies. With *Moirai* we showed that despite the impact of the embedding on the pseudotime prediction, this did not interfere with the interpretation of the data as demonstrated by the great overlap and similarity between the expression patterns recovered. This may motivate the use of *Moirai*, while highlighting the nuance with which pseudotime prediction should be interpreted.

Due to its unique architecture, use of *Moirai* may allow for novel insights into various developmental processes. The identification and use of gene expression patterns during the inference process not only provides a level of internal validation to the method but furthermore eases downstream analyses. It negates the need for clustering and differential expression analysis, which are often employed by other trajectory inference methods, as instead relevant genes are directly inferred based on their expression dynamics across the developmental continuum. Divergent dynamics are also easily derived from *Moirai*’s output and can improve our understanding of how various genes contribute to lineage specification. Future extensions may further enrich *Moirai*’s output and its ability to provide mechanistic insights into the developmental system under study. Altogether, we hope *Moirai* may serve as a useful tool for studying cell differentiation and simultaneously illustrate the value of focusing on small-scale transcriptional events for trajectory inference.

## Data availability

This study did not generate new data; all datasets analysed are publicly available. The human myelopoiesis data (4,142 cells, 16,106 genes) were obtained from the Palantir repository (https://github.com/dpeerlab/Palantir) [27]. The mouse spermatogenesis data (1,829 cells, 54,144 genes), originally generated by Hermann et al. [23], were obtained in processed form from Weiler and Theis [22] via figshare (https://doi.org/10.6084/m9.figshare.28723310). The zebrafish neural crest data (697 cells, 8,012 genes) were obtained from the RegVelo repository (https://github.com/theislab/regvelo) [24]. The acoel *Hofstenia miamia* data (8,405 cells, 20,935 genes) were obtained from https://github.com/JulianKimura/Hulett_etal [25].

## Code availability

*Moirai* is implemented in Python and is available at https://github.com/ jeuken/Moirai.

