## Supplementary information for "Moirai: single-cell trajectory inference grounded in gene-level expression dynamics"

Amber H.B. Fijn [1] and Gustavo S. Jeuken [1,*]

[1]Systems Biology Lab, AIMMS/A-LIFE, Vrije Universiteit Amsterdam, Amsterdam, the Netherlands

[*]Corresponding:

#### Supplementary Note

Though *Moirai* has multiple hyperparameters, a subset of these is most relevant for a successful implementation. We provide some guidance on how these should be tuned here:

- $t_{a_1}$ and $t_{a_2}$: The two autocorrelation thresholds, used for scoring the individual (Methods, Step 5.1) and the aligned (Methods, Step 5.3) random walks, respectively. In this report, values around 0.8 were typically selected because these gave a good tradeoff between quality of the gene expression patterns and the number of gene expression patterns returned. It is typically suitable to set the value of $t_{a_2}$ slightly lower than that of $t_{a_1}$, because it is unlikely that aggregated dynamics achieve a similar autocorrelation score as the individual dynamics.

- $k$: The number of nearest neighbors of the k-NN graph (Methods, Step 1). As discussed in the Methods, the appropriate number of neighbors depends on the density with which cells are represented in the dataset. In larger datasets, it can be expected that cells are relatively more similar to their neighbors, allowing for a relatively higher value of $k$ ($> 20$). For smaller datasets, or datasets where cells are sampled across different timepoints, a smaller value of $k$ ($< 10$) is more appropriate.

- $m$: The maximum number of steps taken by a random walk (Methods, Step 3). The value of this hyperparameter depends on the number of cells in the dataset and the number of neighbors $k$ selected for the transition matrix. Due to random walks being terminated when cycling occurs (i.e., when any cell has been visited more than $t_c$ times during a single random walk), setting $m$ too high will not affect *Moirai*'s output. However, it still comes at a computational cost, underscoring the importance of assessing an appropriate value prior to submitting *Moirai*.

- $h$: The maximum shift applied when aligning a random walk to its seed (Methods, Step 5.3). The value of this hyperparameter depends on the maximum length of the random walks and how starting cells are selected. In general, $h$ cannot be higher than $m$, because this would entail placing two random walks sequentially. Due to the way random walks are initiated, such an expression pattern cannot represent a feature of the developmental process. If the starting population is more widely spread across the graph, a higher $h$ should be selected to allow for alignment of random walks between the population of walks initiated from the different starting cells.

### Supplementary Figures

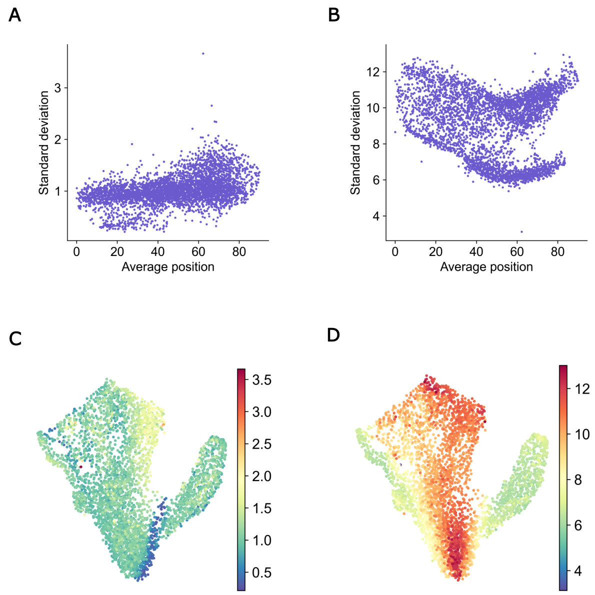


Figure S1: **Standard deviation over pseudotime prediction for human myelopoiesis data**. (**A**) Standard deviation of pseudotime prediction over four iterations with different selections of random walks as guides (Methods, Step 7: Predict pseudotime). Average position refers to the average position of cells in the final alignment matrix $A_f$. (**B**) Standard deviation over position of cells in $A_f$, giving some indication of the uncertainty of a cell's position as inferred by the different gene expression patterns. (**C**) UMAP of myelopoiesis data colored by standard deviation as shown in subplot A. (**D**) UMAP of myelopoiesis data colored by standard deviation as shown in subplot B.

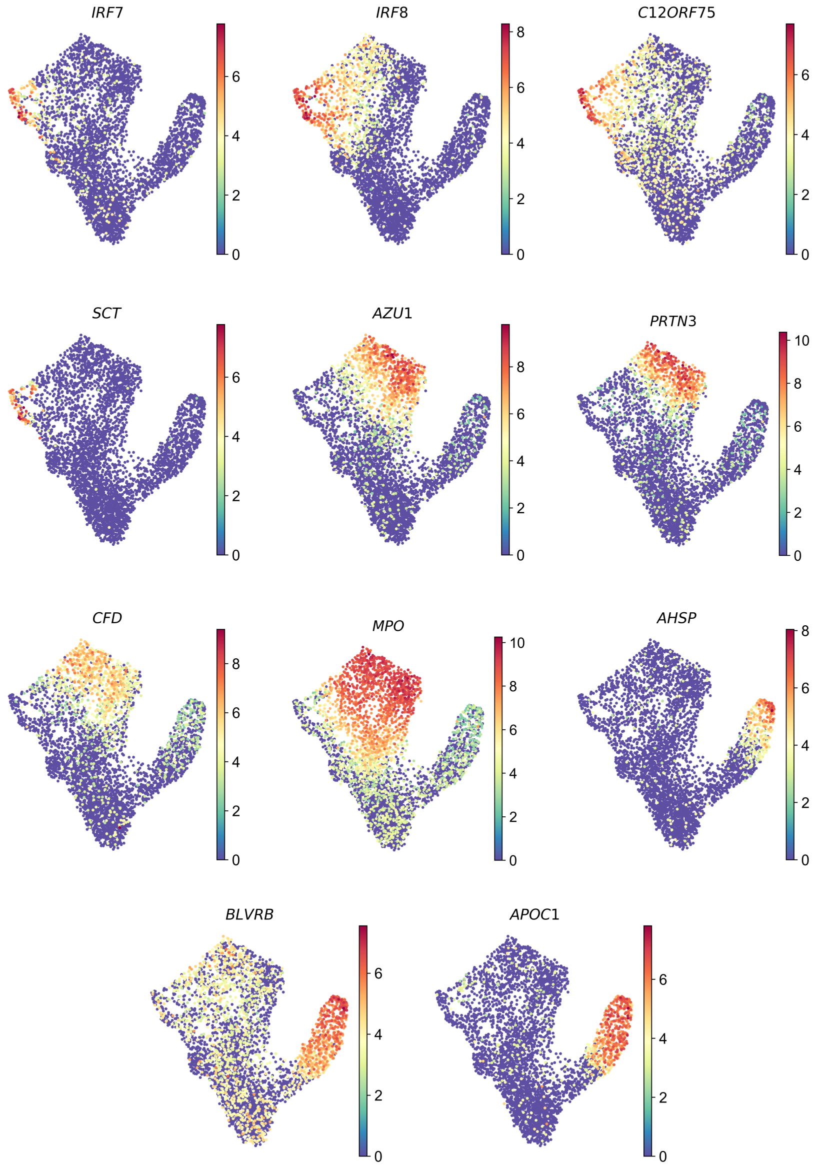


Figure S2: **UMAP of myelopoiesis data colored by expression of 11 genes**. Expression patterns show bias towards distinct lineages.

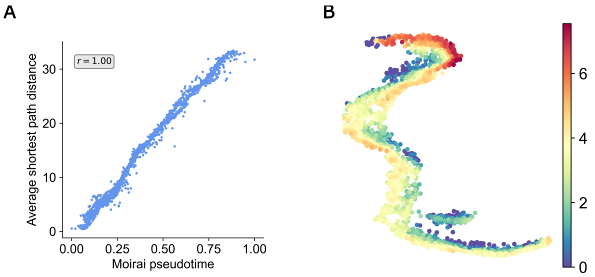


Figure S3: **Additional results from the analysis of the spermatogenesis data**. (**A**) Correlation plot of *Moirai* pseudotime against initial shortest path distance used to construct the transition matrix. Pearson correlation is shown. (**B**) UMAP colored by log visit-frequency of cells by random walks initialized from a population of starting cells.

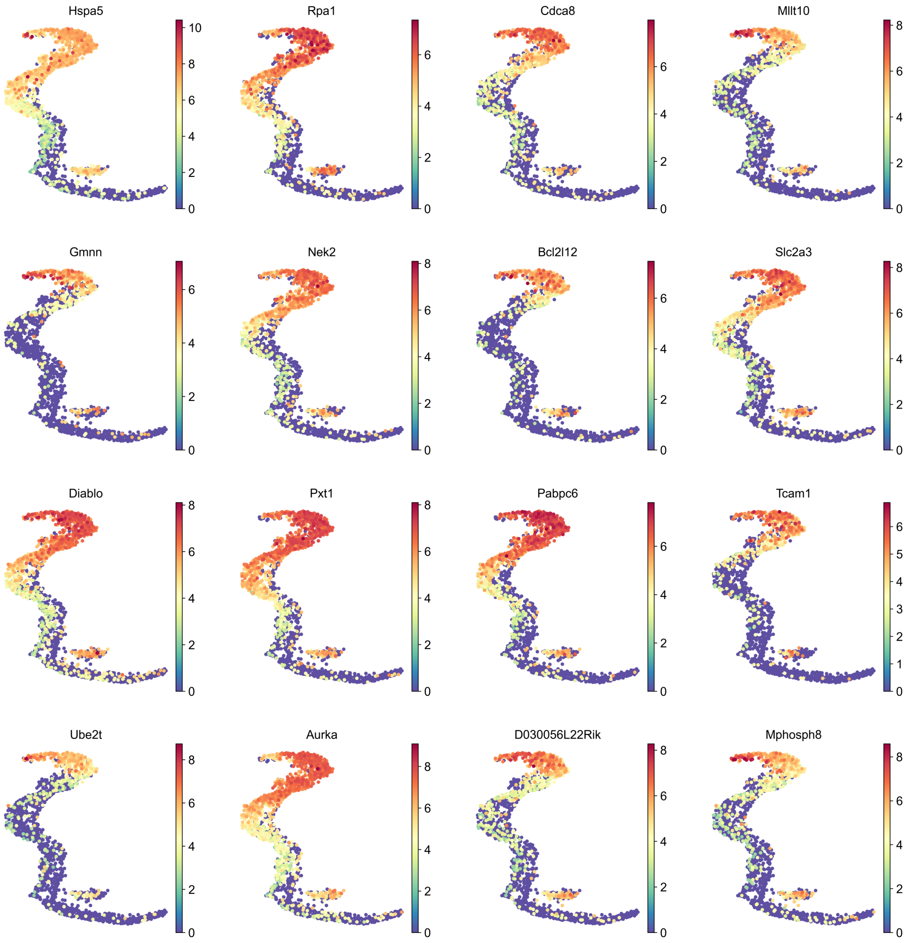


Figure S4: **Distinct population of round spermatid cells shows higher expression magnitude of genes downregulated during spermatogenesis.** UMAP representation colored by expression.

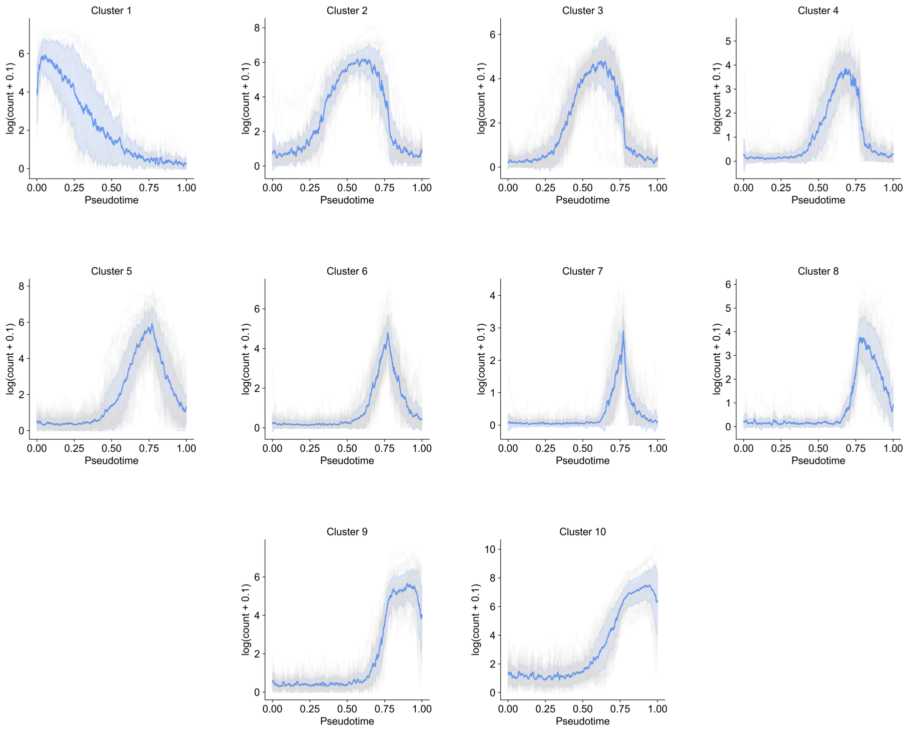


Figure S5: **Cluster analysis reveals ten clusters of genes with distinct expression patterns over spermatogenesis pseudotime**. Cluster analysis was performed for those genes identified by *Moirai* during the inference process. For both analysis and visualization, moving average was used.

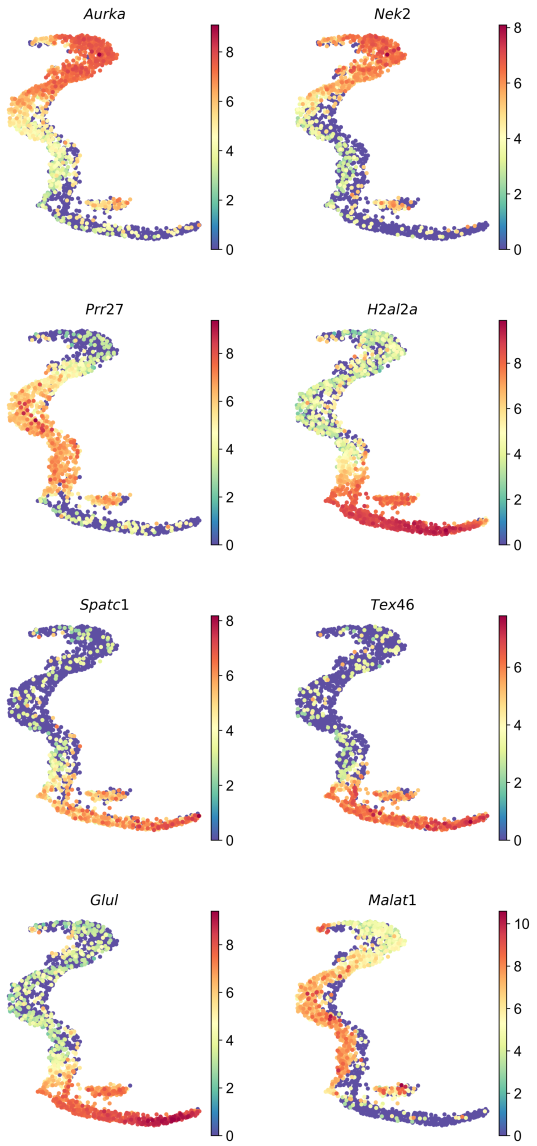


Figure S6: **UMAP of spermatogenesis data colored by expression of 11 genes**. The genes are upregulated during distinct windows of spermatogenesis.

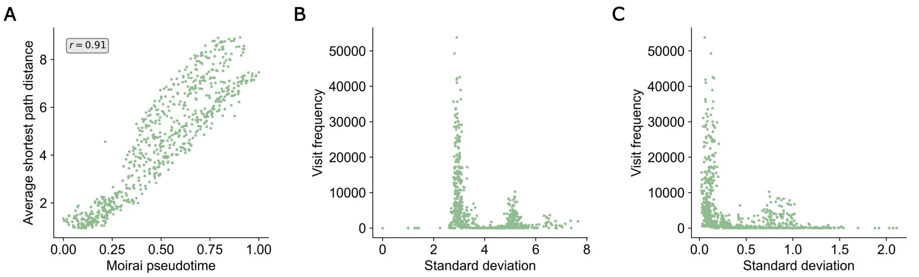


Figure S7: **Additional results from the analysis of the zebrafish data**. (**A**) Correlation plot of *Moirai* pseudotime against initial shortest path distance used to construct the transition matrix. Pearson correlation is shown. (**B**) Scatter plot showing relation between the amount of times a cell is represented in the final alignment matrix $A_f$ and the standard deviation of that cell's pseudotime over four iterations of running pseudotime prediction with different random walks acting as guides (Methods, Step 7: Predict pseudotime). (**C**) Scatter plot showing relation between the amount of times a cell is represented in $A_f$ and the standard deviation over the position of that cell in $A_f$.

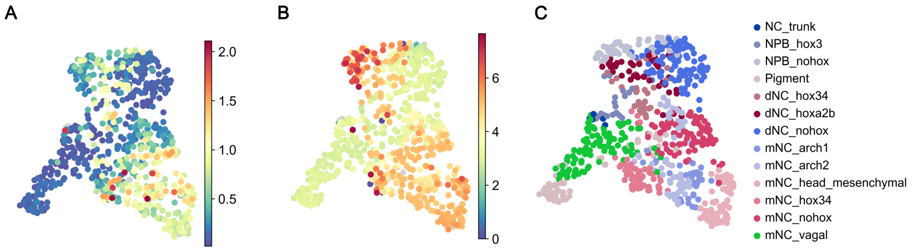


Figure S8: **Standard deviation over pseudotime prediction for zebrafish neural crest data**. (**A**) UMAP colored by standard deviation of pseudotime prediction over four iterations with different selections of random walks as guides (Methods, Step 7: Predict pseudotime). (**B**) UMAP colored by standard deviation over position of cells in $A_f$. (**C**) UMAP with cell type annotation from Wang et al. [1].

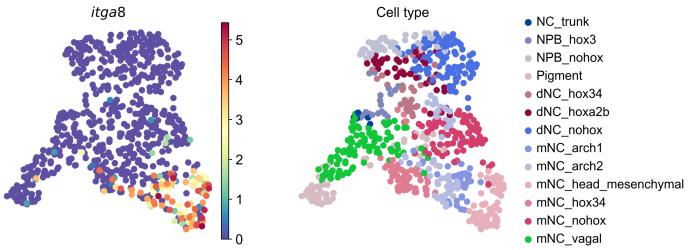


Figure S9: **Expression of *itga8* is biased towards cells of the pharyngeal arches**. On the left, the UMAP of the zebrafish neural crest data is colored by *itga8* expression. The right plot shows the original annotation from Wang et al. [1].

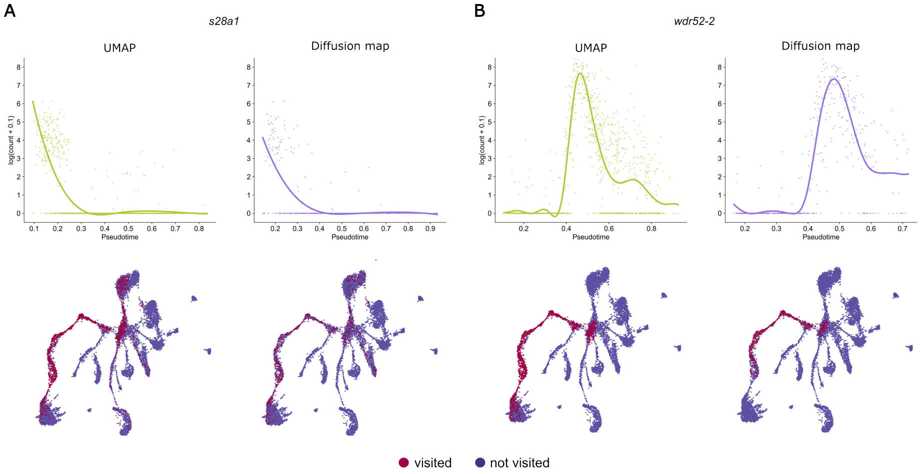


Figure S10: **Expression pattern and random walk coverage of *s28a1* and *wdr52-2* for *Moirai* output derived from two embeddings**. (**A**) Expression pattern and coverage of random walks of *s28a1* for the UMAP and diffusion map derived results. Coverage refers to the cells visited as part of the random walks that captured smooth expression patterns for *s28a1*. (**B**) Expression pattern and coverage of *wdr52-2* captured for the UMAP and diffusion map derived results.

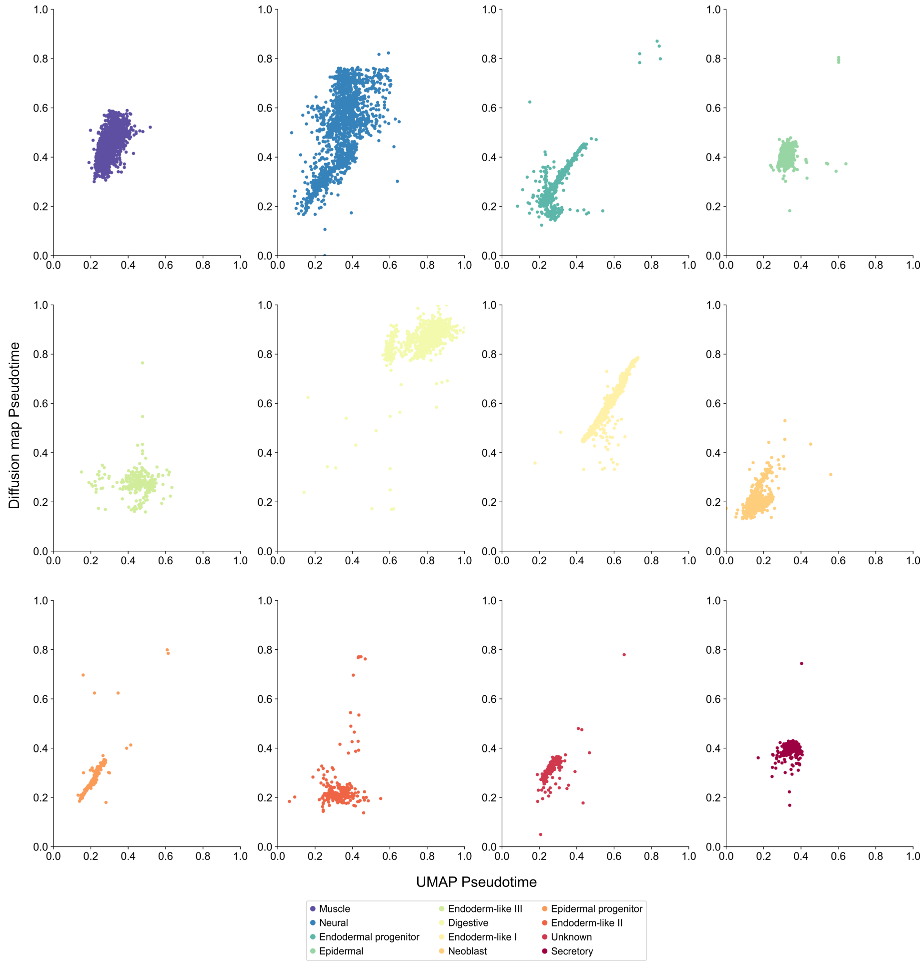


Figure S11: **Correlation plots of *Moirai*'s pseudotime predicted using UMAP and diffusion map embeddings for acoel data**. Subplots display the correlation for each cell type of the acoel embryogenesis data.

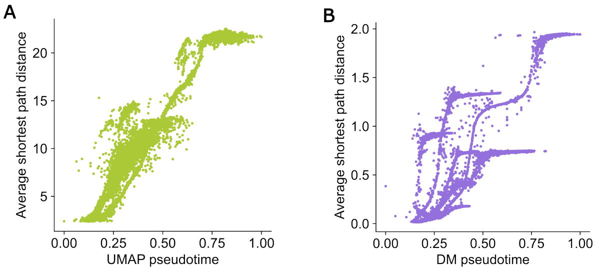


Figure S12: **Correlation of final pseudotime prediction with shortest-path distance of the acoel data**. Shown for the (**A**) UMAP and (**B**) diffusion map derived embeddings.

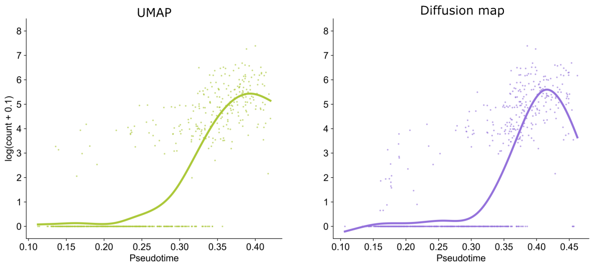


Figure S13: **Expression pattern of *tenx-3* captured for the acoel data**. The left panel shows the expression pattern captured from the UMAP embedding and the right that for the diffusion map embedding.

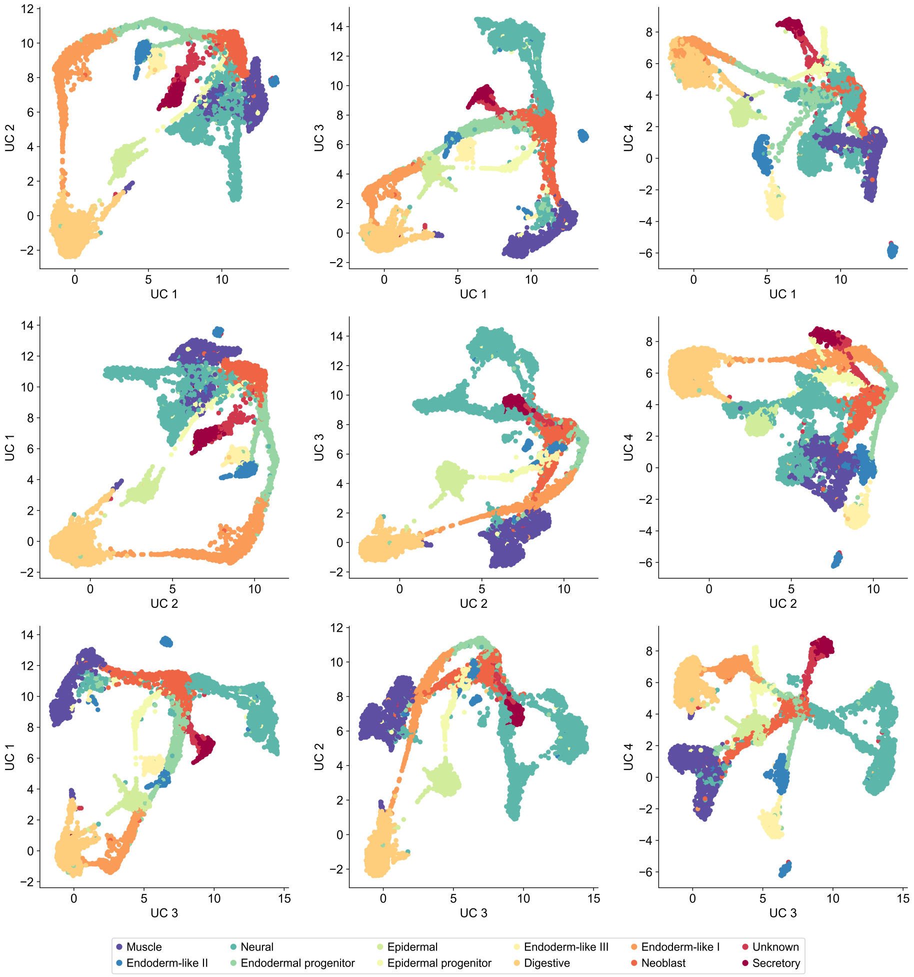


Figure S14: **UMAP of acoel data**. The four-dimensional embedding is shown for every unique projection.

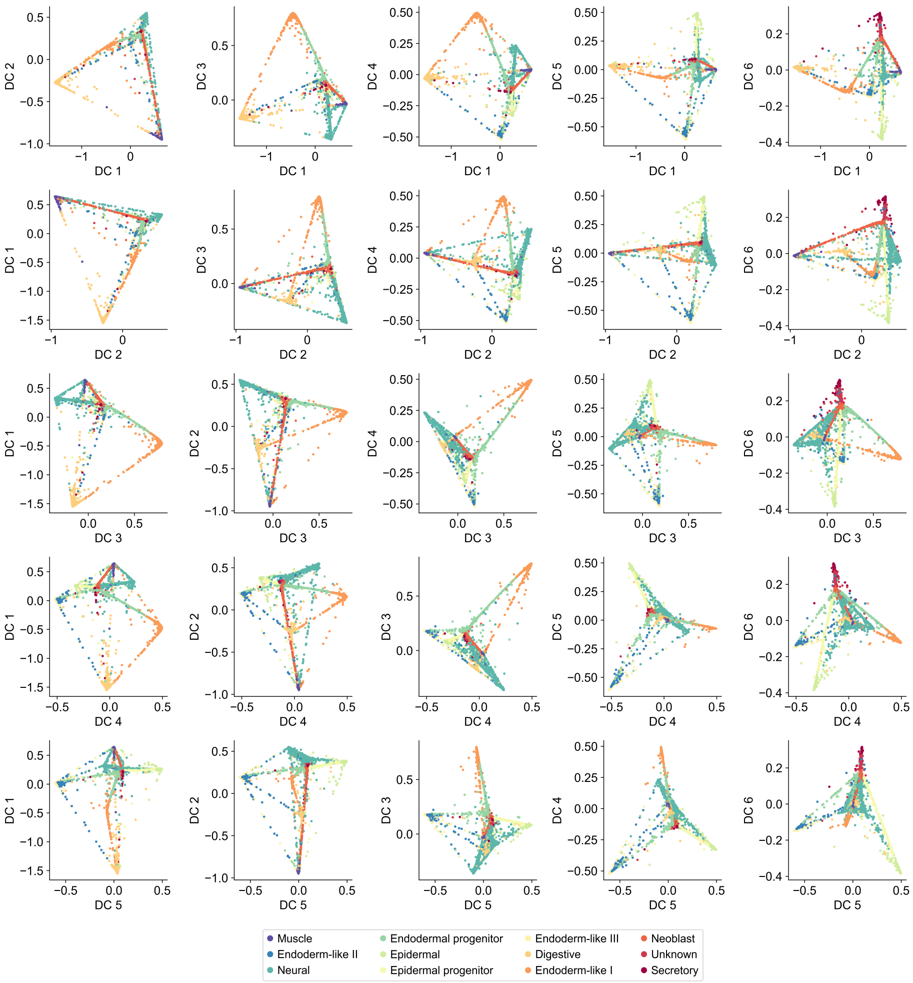


Figure S15: **Diffusion map of acoel data**. The six-dimensional embedding is shown from every unique projection.
